# Fibroblast Growth Factor Receptor Signaling in Maturing Osteoblasts Controls Cell-Matrix Interactions Critical for Osteocyte Survival

**DOI:** 10.64898/2026.04.27.721119

**Authors:** Debabrata Patra, Craig Smith, Chenming Wei, Courtney M. Mazur, Ilemona Ameadaji, Tiandao Li, Marc N. Wein, Mathew J. Silva, David M. Ornitz

## Abstract

The terminal differentiation of osteoblasts into osteocytes, the most abundant cell type in cortical bone, is critical for skeletal homeostasis. Osteocyte loss is a hallmark of bone aging and fragility, yet the mechanisms regulating osteocyte formation and survival are poorly understood. We show that inactivation of fibroblast growth factor receptor 1 (*Fgfr1*) in the mature osteoblast lineage results in extensive osteocyte death, identifying FGFR1 signaling as essential for osteocyte viability and bone integrity. Lineage tracing and analysis of endogenous and induced appositional bone formation revealed that newly embedded osteocytes fail to survive without FGFR1. These osteocytes exhibited ectopic expression of osteocalcin and podoplanin within sclerostin-positive, TUNEL-reactive lacunae, along with defective dendrite formation and disruption of the local lacunocanalicular network. RNA sequencing of cortical bone demonstrated reduced expression of extracellular matrix (ECM) genes and neuronal regulatory genes, while histological and ultrastructural analyses showed disorganized collagen fibrils, diminished osteoid, and abnormal mineralization. In vitro, FGF signaling in Ocy454 cells regulated gene programs involved in development, axon guidance, and bone ECM organization, highlighting a dual function for FGF signaling in which it controls both matrix-dependent and intrinsic cell differentiation mechanisms during the osteoblast-to-osteocyte transition. We propose that FGFR1 deficiency causes ECM disorganization and impaired dendrite formation, disrupting osteocyte communication with neighboring bone and vascular cells, ultimately leading to cell death. These findings establish FGFR signaling as a critical regulator of osteocyte differentiation, viability of bone-embedded osteocytes, and bone homeostasis.

**Summary Statement:** FGFR signaling has a profound effect on adult bone extracellular matrix that is vital to maintaining the viability and morphology of newly formed osteocytes, their lacunocanalicular network and the maintenance of bone homeostasis.

## Introduction

The osteocyte is the predominant cell type of the mammalian skeleton and they are the most differentiated cell type present in bone at any given stage ^1^. With a half-life of several years to even decades ^2^ they are long-lived cells and a major focus in osteocyte biology is to correlate their viability, longevity and functions with skeletal and overall health in the aging population ^3, 4^.

Functionally, osteocytes are important for bone homeostasis. They are master regulators that control osteoblast-directed bone formation and osteoclast-directed bone resorption ^5^. They regulate skeletal remodeling via secretion of sclerostin (SOST) and Dickkopf-1 (DKK1) which are negative regulators of anabolic Wnt signaling in osteoblasts, and they secrete Receptor Activator of Nuclear Factor Kappa-B Ligand (RANKL), which modulates osteoclast formation and function ^6, 7^. Osteocytes also function as an endocrine organ, secreting FGF23 which regulates mineralization ^8^ and phosphate homeostasis ^9^. A characteristic and important functional feature of the osteocyte is their ability to make cellular projections called dendrites that form the extensive, interconnected lacunocanalicular network (LCN), which traverses the entire length of a bone ^10^. The uniform distribution of osteocytes in cortical bone and their connectivity to each other and to other cells associated with the skeletal system, such as bone lining cells, osteoblasts and vascular cells via the LCN network and gap junctions provide osteocytes with the unique ability to be the primary sensor cell that regulates skeletal remodeling.

The mechanisms regulating the differentiation of the mature osteoblast to an osteocyte are not completely understood but begins with the embedding of the osteoblast in the osteoid, the collagenous, non-mineralized matrix, in newly formed bone. Concomitant to embedding, differentiating osteocytes also generate dendrites that will eventually form the LCN ^11^. Mineralization of the osteoid results in the formation of the mineralized bone matrix which encapsulates the osteocyte in a lacuna where it remains embedded for its lifetime, functioning as an active multifunctional cell in conjunction with the LCN. This differentiation process results in the loss of osteoblast-related markers such as osteocalcin (OCN, *Bglap*) and Type I Collagen (Col1a1) and the gain of osteocyte-specific markers, chief among them sclerostin (SOST), which marks the complete differentiation of an osteoblast to an osteocyte ^2, 10^. Early markers of this differentiation process, such as dentin matrix protein (DMP1) and podoplanin (PDPN/E11/gp38) mark the initiation of the osteoblast to osteocyte transition ^2, 12, 13^.

Fibroblast growth factor (FGF) signaling is vital to skeletal development with established functions in both chondrogenesis and osteogenesis ^14–16^. The critical role of FGF signaling in skeletal development was established when mutations in humans in FGF receptor 1 (FGFR1), FGFR2 and FGFR3 were discovered to result in ligand-independent activation of these receptors resulting in Achondroplasia, Crouzon, Apert, Pfeifer and bent bone dysplasia syndromes ^17^. A loss of function mutation in FGFR3 has also been documented in the CATSHL syndrome that results in skeletal overgrowth due to increased chondrogenesis. Mutations in FGF/FGFR signaling that result in skeletal deformities are not unexpected given the expression of FGFRs in most chondrogenic and osteogenic cell lineages not only during development but also in the postnatal and mature bone. In the mature bone, FGFR1 expression is seen in the osteoblasts, in bone lining cells, and in the osteocytes of cortical and trabecular bone. Postnatally, FGFR1 is also expressed in the prehypertrophic and hypertrophic chondrocytes in the growth plate ^17–20^.

Mechanistically the function(s) of FGFR1 and FGFR2 in mature osteolineage cells is not known. In a previous study, both *Fgfr1* and *Fgfr2* were inactivated in mature osteoblasts and osteocytes in growing bone via osteocalcin-Cre (OCN-Cre), which becomes active at about embryonic day 17, or with Dentin Matrix Protein 1 (DMP1)-CreER in juvenile mice. In both cases, significant osteocyte death was observed accompanied by a striking increase in local bone mass after 6-12 weeks ^21^. Gene expression studies suggested that loss of sclerostin and activation of Wnt/β-catenin signaling could contribute to the increased bone mass.

To determine if loss of FGFR signaling affects osteocyte viability independent of longitudinal bone growth, we conditionally inactivated *Fgfr1* and *Fgfr2* in 12-week-old mice. Here, we show that inactivation of *Fgfr1* in adult bone leads to osteocyte death and that abnormal bone phenotypes are limited to regions of new appositional bone growth indicating a mechanistic link to new bone formation. Gene expression analysis *in vitro* and *in vivo* showed that FGFR signaling regulated dendritic outgrowth and genes involved in neuronal development, axon guidance and formation of bone ECM. These observations establish the importance of FGFR signaling in the regulation of osteoblast to osteocyte transition, and mechanistically in the deposition and organization of the bone ECM and the maturation and survival of bone-embedded osteocytes.

## Results

### DMP1-CreER directed *Fgfr1* and *Fgfr2* inactivation results in osteocyte apoptosis in adult mice

To determine the role of FGFR signaling in mature adult bone without influence from development, the inducible DMP1-CreER allele was used to inactivate *Fgfr1* and/or *Fgfr2* in adult (12-week-old) mice. DMP1-CreER targets the mature osteoblast and osteocyte lineage with minimal activity detected in non-skeletal tissue. On exposure to tamoxifen, DMP1-CreER-dependent reporter expression (*ROSA^TDT^*) confirmed Cre activity only in the endosteum, periosteum and osteocytes (Fig. 1a). Floxed alleles of *Fgfr1* or *Fgfr1* and *Fgfr2* (DFF) were bred with DMP1-CreER transgenic mice to generate DMP1-CreER; *Fgfr1^f/f^* conditional knockout (FGFR1-CKO) or DMP1-CreER; *Fgfr1^f/f^; Fgfr2^f/f^*double conditional knockout (FGFR1,2-DCKO) mice. Twelve-week-old Control and FGFR1,2-DCKO mice were induced with tamoxifen for four weeks (Fig. 1b). Quantitative RT-PCR (qRT-PCR) using mRNA made from flushed cortical bones confirmed reduced *Fgfr1* and *Fgfr2* levels in FGFR1,*2*-DCKO compared to Control DFF mice (Fig. 1c). FGFR1-CKO mice that were not exposed to tamoxifen demonstrated low background activity from the *ROSA^TDT^* reporter allele (Fig. 1d).

**Figure 1.**
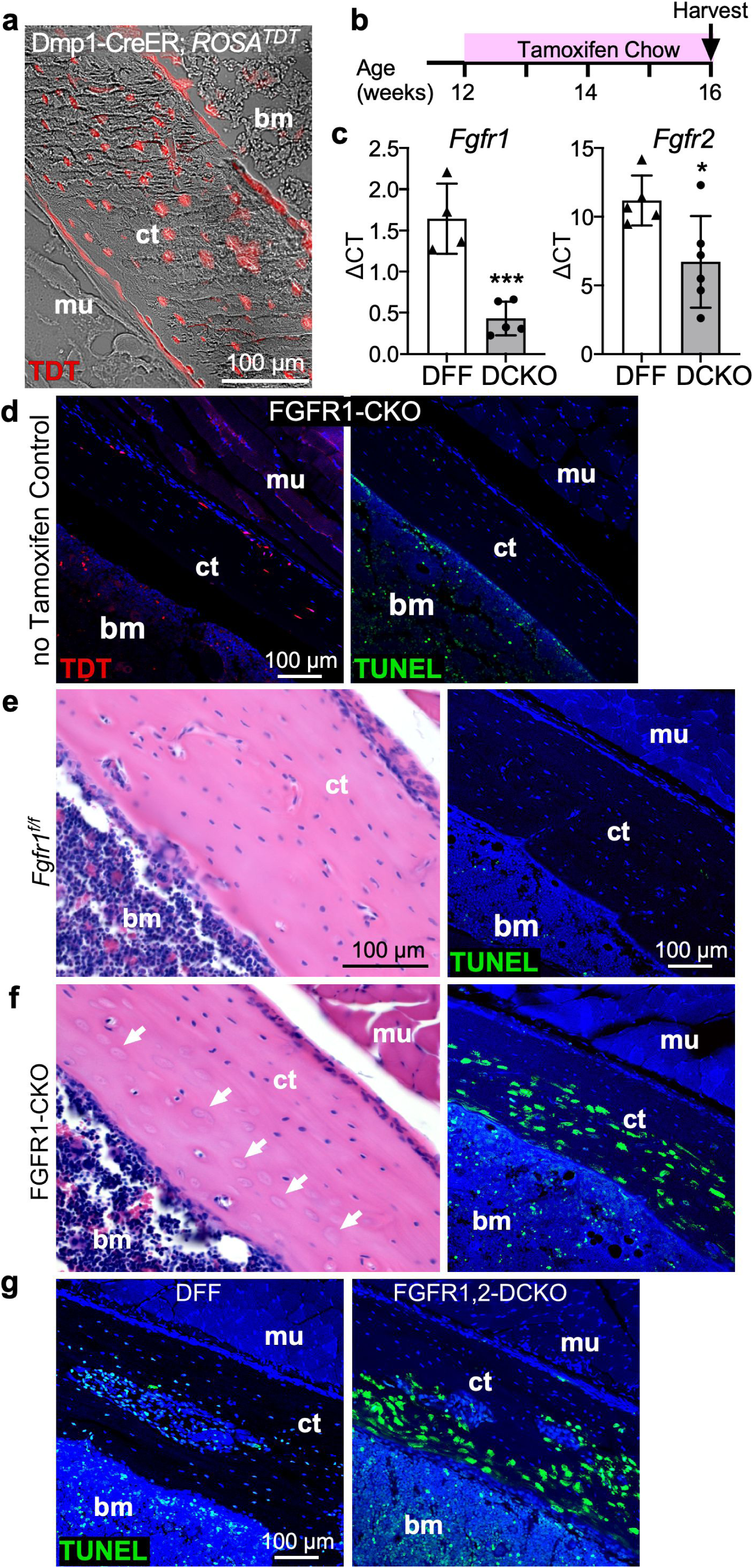
*Fgfr1* and *Fgfr1, 2* inactivation in the DMP1 lineage in adult mice results in osteocyte death by apoptosis. **(a)** DMP1-CreER; *ROSA^TDT^* mice were treated with tamoxifen from 3 to 5 weeks of age and bones harvested at 6 weeks of age. The DMP1-CreER targeted lineage (TDT expression) marks osteoblasts in the periosteum, endosteum and osteocytes within cortical bone. **(b)** Experimental plan in which 12-week-old mice were exposed to tamoxifen for 4 weeks to induce DMP1-CreER. **(c)** Quantitative RT-PCR analysis of *Fgfr1* and *Fgfr2* from mRNA harvested from the femora and tibiae of *Fgfr1^f/f^; Fgfr2^f/f^* (DFF) and DMP1-CreER; *Fgfr1^f/f^*; *Fgfr2^f/f^*(FGFR1,2-DCKO) cortical bones. Data shown as the mean ± SD. Unpaired *t*-test, *n*=4-5, \**P*<0.05, \*\*\**P*<0.001. **(d)** IF for *ROSA^TDT^*(TDT) reporter (indicator of DMP1-CreER induction; red) and TUNEL assay in the tibia of FGFR1-CKO mice without exposure to tamoxifen. (**e, f**) Hind limbs were harvested after four weeks of tamoxifen exposure from 16-week-old *Fgfr1^f/f^* Control (**e**) and DMP1-CreER; *Fgfr1^f/f^*(FGFR1-CKO) (**f**) mice. Tibiae were analyzed by H&E staining (left) and TUNEL assay (green) to assess apoptosis (right). Arrows in (**f**) show the absence of nuclear staining in the lacunae in FGFR1-CKO mice. **(g)** TUNEL assay comparing DFF Control and FGFR1,2-DCKO mice. Blue; DAPI-stained nuclei; bm, bone marrow; ct, cortical bone; mu, muscle. Scale bars: 100 µm.

Hindlimbs from 16-week-old mice were analyzed for histological changes and cell death. Like OCN-Cre targeted inactivation of *Fgfr1* and *Fgfr2* ^21^, DMP1-CreER inactivation of *Fgfr1* (FGFR1-CKO) (Fig. 1e, f; Fig. S1a) or both *Fgfr1* and *Fgfr2* (FGFR1,2-DCKO) (Fig. 1g) resulted in osteocyte death by apoptosis as seen by empty cortical bone lacunae and TUNEL positive lacunae in conditional knockout but not in Control mice. Both male and female CKO/DCKO mice demonstrated TUNEL positive osteocytes (7 male, 9 female), compared to control mice (10 male, 11 female) which were TUNEL negative. Additionally, FGFR1-CKO mice that did not receive tamoxifen showed no TUNEL+ osteocytes (Fig. 1d) indicating that osteocyte cell death on exposure to tamoxifen was specifically due to inactivation of FGFR signaling.

Interestingly both H&E and TUNEL staining revealed that dying osteocytes are localized closer to the endosteum in the mid- and distal-diaphyseal regions, while osteocytes closer to the periosteum remain unaffected (Fig. 1e-g; Fig. S1a). Our previous study suggested that loss of *Fgfr1*, and not *Fgfr2*, is responsible for osteocyte apoptosis ^21^, which is consistent with similar phenotypes in FGFR1-CKO and FGFR1,2-DCKO mice. To determine if inactivation of *Fgfr2* in adult mouse osteoblasts affect osteocyte viability, we created DMP1-CreER; *Fgfr2^f/f^* (FGFR2-CKO) mice. FGFR2-CKO mice were induced with tamoxifen from 12-16 weeks of age. Histological analysis of tibia from 16-week-old mice showed no evidence of empty lacunae or TUNEL positive lacunae in cortical bone (Fig. S2). In subsequent experiments we have used FGFR1-CKO and FGFR1,2-DCKO mice interchangeably for our studies.

### Abnormal markers of osteoblast and osteocyte differentiation in the zone of TUNEL+ cells

To investigate the mechanism of osteocyte apoptosis triggered by *Fgfr* inactivation in adult bones, immunofluorescence (IF) staining of cortical bone was used to assess expression of osteoblast and osteocyte proteins. Sclerostin (SOST) is expressed in mature osteocytes but not osteoblasts ^22^. IF for SOST showed similar expression patterns in both Control and FGFR1-CKO osteocytes indicative of osteoblast to osteocyte differentiation even in the absence of FGFR1 signaling (Fig. 2a). These mice did not show expression of osterix (Osx, a pre-osteoblast marker) in cortical bone osteocytes (Fig. 2b), even though its expression was observed in the growth plate and in the trabecular osteoblasts, as expected (Fig. 2c). Likewise, periostin (POSTN, a periosteal marker) expression was observed only in the periosteum, but not in the osteocytes (Fig. S3a). However, IF analysis for osteocalcin (OCN, *Bglap*), a mature osteoblast marker ^10, 23^ demonstrated abnormally high expression in FGFR1-CKO osteocytes (Fig. 2f; Fig. S1b). Remarkably, OCN expression was observed only in osteocytes present within the zone of osteocyte cell death, closer to the endosteum; OCN expression was absent in osteocytes closer to the periosteum where apoptosis was not observed (Fig. S1a, b). Next, we performed IF for podoplanin (PDPN, E11/T1a), a protein expressed by osteocytes in the early stages of differentiation but not in mature osteocytes ^10, 12, 13^. Abnormal PDPN/E11 expression was seen in FGFR1-CKO osteocytes but not in Control osteocytes (Fig. 2g; Fig. S1c), mirroring the expression pattern of OCN expression in the zone of TUNEL+ osteocytes (Fig. S1a, b). Double-labeled IF analyses for both OCN (green) and PDPN (red) showed the co-expression of these proteins (yellow) in many osteocytes within this zone (Fig. 2h). Like OCN, osteocytes in this zone also showed expression of alkaline phosphatase (ALPL), a mature osteoblast, but not osteocyte, marker (Fig. 2i) ^23^. The presence of OCN-expressing (not shown), PDPN-expressing and ALPL-expressing osteocytes mirroring the location of TUNEL+ osteocytes was also seen in FGFR1,2-DCKO mice (Fig. S3b, c). Analysis of both TUNEL and OCN (Fig. S4a-c) or TUNEL and ALPL (Fig. S4d-f) indicated that TUNEL+ osteocytes also express OCN and ALPL. Taken together, these observations suggest that *Fgfr1*inactivation results in abnormal osteoblast to osteocyte differentiation with retention of osteoblast markers.

**Figure 2.**
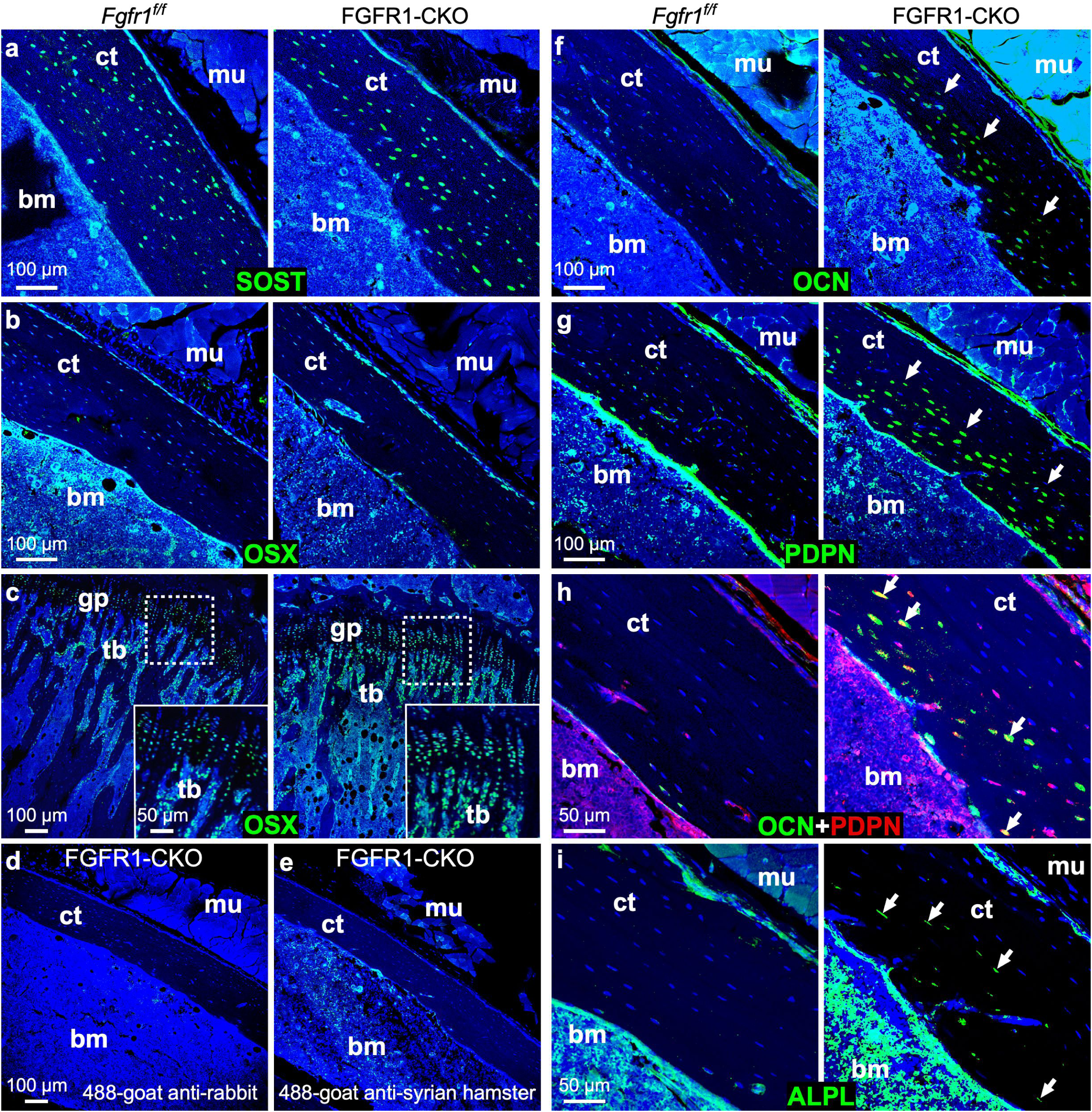
*Fgfr1* inactivation in the DMP1 lineage results in abnormal osteocyte phenotypes. Mice were exposed to tamoxifen from 12-16 weeks of age and tibiae were harvested for IF. (**a**) IF for sclerostin (SOST) (green) showing uniform labeling throughout cortical bone. (**b, c**) IF for osterix (Osx) (green) showing absence of signal within adult cortical bone (**b**) but expected signals observed in the growth plate and in trabecular osteoblasts (**c**) in both Control and FGFR1-CKO mice. (**d, e**) Negative Controls for IF with no primary antibody shown for FGFR1-CKO tibiae stained with 488-goat anti-rabbit (**d**, secondary antibody for OCN) or 488-goat anti-Syrian hamster (**e**, secondary antibody for PDPN). (**f, g**) IF for osteocalcin (OCN) (green) (**f**) and podoplanin (PDPN) (green) (**g**). Arrows indicate staining on the endosteal side of the tibia. **(h)** Double labeled IF for OCN (red) and PDPN (green). Arrows indicate osteocytes in the endosteal region of tibia that stain for both markers (yellow). **(i)** IF for bone-specific alkaline phosphatase (ALPL) (green). Arrows indicate staining on the endosteal side of the tibia. Blue, DAPI-stained nuclei; bm, bone marrow; ct, cortical bone; gp, growth plate; tb, trabecular bone. Scale bars: (a-g) 100 µm; (h,i) 50 µm.

To determine whether abnormal gene expression precedes the onset of TUNEL+ cells, we performed a temporal analysis (Fig. 3). Control (*Fgfr1^f/f^*) and FGFR1-CKO mice were exposed to tamoxifen chow beginning at 12 weeks of age and harvested at one-week intervals for TUNEL, *ROSA^TDT^* reporter, PDPN and OCN analyses. As expected, osteocytes in Control (*Fgfr1^f/f^*) mice did not show any evidence of apoptosis at any time point (Fig. 3a). At the end of one or two weeks, FGFR1-CKO mice did not show any evidence of apoptosis (Fig. 3b), nor did they demonstrate any ectopic expression of PDPN and OCN (Fig. 3d, e) even though significant expression of *ROSA^TDT^* was observed (Fig. 3c), indicating sufficient DMP1-CreER induction in the osteoblast-osteocyte lineage. However, three weeks after induction the first signs of TUNEL+ osteocytes were observed, concurrent with ectopic PDPN and OCN expression, suggesting that at least three weeks of DMP1-CreER induction is required to observe osteocyte apoptosis. These observations suggest that abnormal expressions of PDPN and OCN are unlikely to be a trigger for inducing osteocyte apoptosis in the absence of FGFR1.

**Figure 3.**
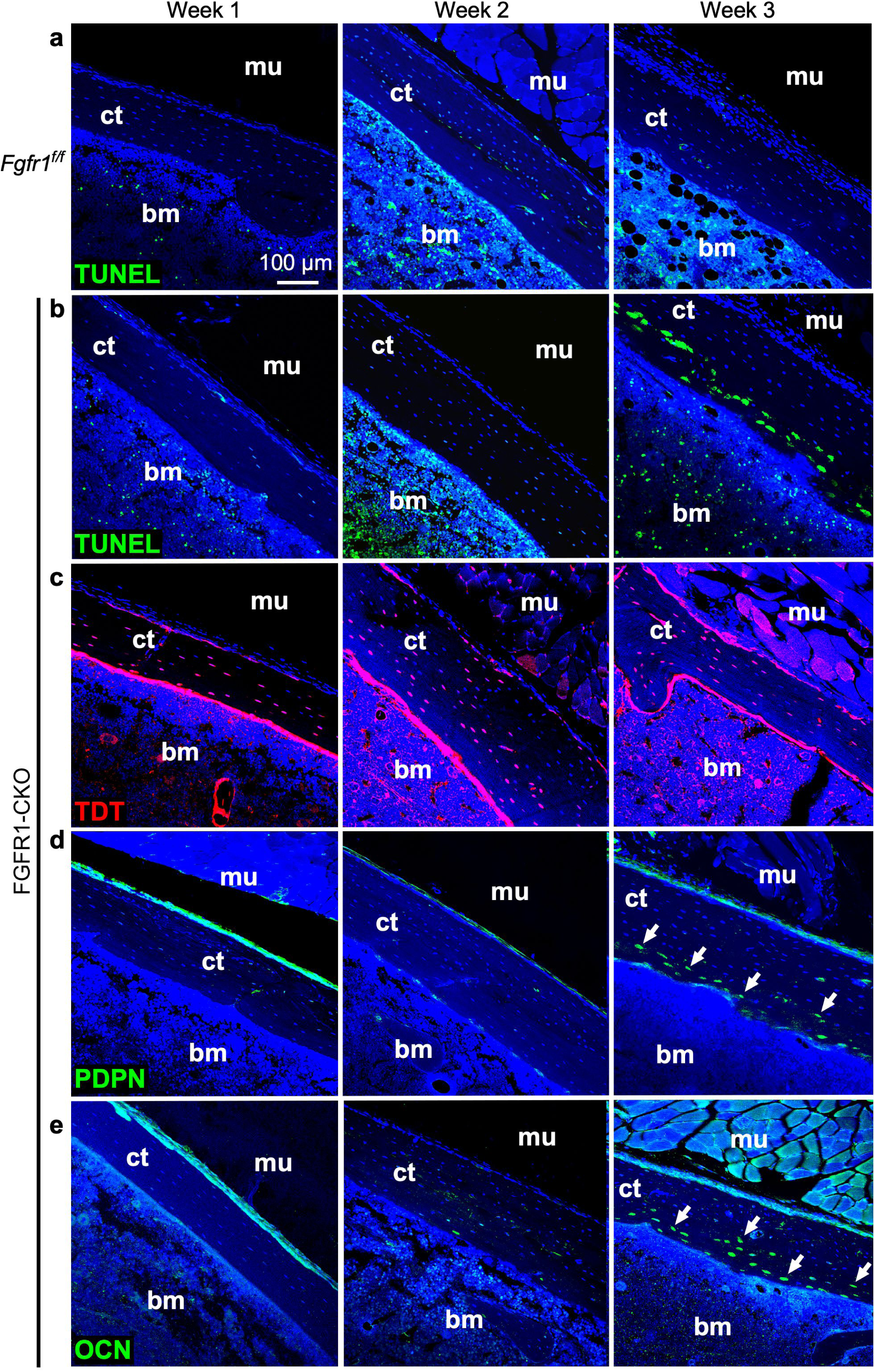
Temporal analysis of apoptosis induction and expression of PDPN and OCN in the FGFR1-CKO cortical bone. Mice were exposed to tamoxifen beginning at 12 weeks of age and tibiae were harvested for analyses after 1, 2 or 3 weeks. **(a)** *Fgfr1^f/f^* Control mice were assessed for apoptosis via TUNEL assay (green). (**b-e**) FGFR1-CKO mice were analyzed for apoptosis (TUNEL; green) (**b**) and IF for *ROSA^TDT^* (TDT) reporter (red) (**c**), PDPN (green) (**d**) and OCN (green) (**e**). Arrows indicate PDPN/OCN expression in the FGFR1-CKO tibiae seen first at 3 weeks post DMP1-CreER induction concurrent to development of TUNEL+ osteocytes. Blue; DAPI-stained nuclei; bm, bone marrow; ct, cortical bone. Scale bars: (all panels) 100 µm.

### Loss of the lacunocanalicular network (LCN) with *Fgfr* inactivation in the DMP1 lineage

The LCN is a characteristic feature of the osteocyte and is central to its functions of mechanosensing and communication with other osteocytes, osteoblasts, bone marrow, vasculature and other organs ^24^. To determine if *Fgfr* inactivation affects the LCN, we performed silver staining and phalloidin F-actin staining to visualize the LCN in Control, FGFR1-CKO and FGFR1,2-DCKO mice. Following 4 weeks of tamoxifen induction, *Fgfr1* inactivation resulted in a dramatic loss of the LCN in a manner that mirrored the location of TUNEL+ cells (Fig. 4a-c; Fig. S5). Loss of the LCN was seen closer to the endosteum in both the midshaft and distal bone, with regions closer to the periosteum showing neither TUNEL+ osteocytes nor LCN loss in both FGFR1-CKO (Fig. 4a-c; Fig. S5) and FGFR1,2-DCKO mice (Fig. 4d, e). FGFR1-CKO and FGFR1,2-DCKO mice demonstrated a large reduction in the average number of dendrites per osteocyte/mouse on the endosteal side of the cortical bone (Fig. 4f). To investigate if loss of the LCN preceded osteocyte apoptosis, we performed a temporal investigation of the loss of LCN (not shown). Like PDPN and OCN expressions, loss of the LCN also appeared to occur concurrently with the onset of osteocyte apoptosis.

**Figure 4.**
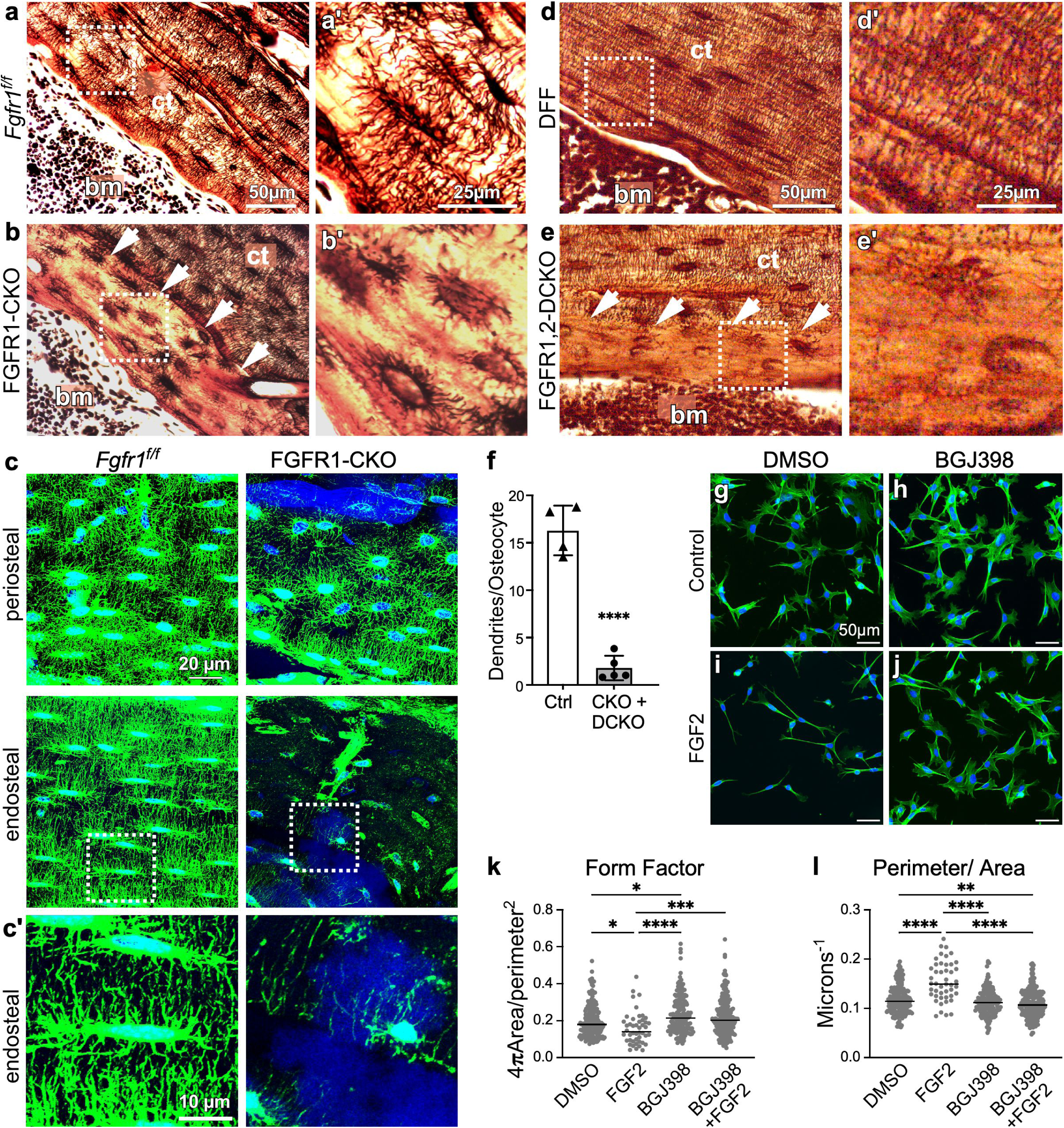
Severe reduction of the lacunocanalicular network (LCN) in the zone of TUNEL+ osteocytes. (**a-c**) Silver staining (brown) and phalloidin staining (green) of the LCN of Control *Fgfr1^f/f^*(**a, c**) and FGFR1-CKO (**b, c**) tibia following induction with tamoxifen from 12 to 16 weeks of age. (**a’, b’, c’**) High magnification images of boxed areas indicated in corresponding panels. (**d, e**) Silver staining (brown) of Control DFF (**d**) and FGFR1,2-DCKO (**e**) tibia following induction with tamoxifen from 12 to 16 weeks of age. (**d’, e’**) High magnification images of boxed areas indicated in corresponding panels. Arrows indicate the edge of the intact LCN within the cortical bone and show a severe reduction in LCN staining between the arrow to the endosteum in FGFR1-CKO and FGFR1,2-DCKO tibiae. Blue; DAPI-stained nuclei; bm, bone marrow; ct, cortical bone. Scale bars: 10, 20, 25 and 50 µm as indicated. (**f**) Average number of dendrites/osteocyte/mouse on the endosteal side of cortical bone in control (n=4) and FGFR1-CKO/FGFR1,2-DCKO (n=5) mice. Only dendrites that were at least 10 µm in length were counted. Data is shown as mean ± SD. Unpaired *t*-test of Control compared to combined CKO plus DCKO data, \*\*\*\**P*<0.0001. (**g-j**) Ocy454 osteocyte-like cells treated for 48 hr with FGF2 (5 ng/mL) (**i, j**) with (**h, j**) or without (**g, i**) BGJ398 (200 nM), visualized with phalloidin (green) and DAPI (blue). Scale bar: 50 µm. (**k, l**) Analysis of the cell shape of Ocy454 cells following 48 hr of the indicated treatments, indicating changes in form factor (**k**) and the perimeter/area ratio (**l**). Each point represents one cell with the line indicating group median. n=46-248 cells. ANOVA with Dunn’s multiple comparisons test, * *P*<0.05, ** *P*<0.01, *** *P*<0.001, **** *P*<0.0001.

### Inhibition of FGFR signaling in Ocy454 osteocyte-like cells supports a role for FGF signaling in dendrite outgrowth and ECM organization

To determine if osteocytes can directly respond to FGF we used the Ocy454 cell line, a conditionally immortalized osteocyte-like cell derived from mouse bone ^25, 26^. Control Ocy454 cells have an actin-rich cytoskeleton with numerous dendritic projections as visualized with phalloidin (Fig 4g). Treatment with the FGFR-inhibitor BGJ398 for 48 hr causes cells to spread out with an enlarged cytoskeletal area and shorter dendritic projections (Fig. 4h). In contrast, cells cultured in the presence of FGF2 for 48 hr adopted an extended morphology with a few long dendrites on each cell (Fig. 4i). BGJ398 treatment blocked these effects of FGF2 on cell morphology (Fig. 4j; Fig. S6a) indicating that BGJ398 is an effective FGFR inhibitor and that the effects of FGF2 are dependent on FGFR signaling in Ocy454 cells. Quantitative analysis of cell shape ^27^ showed a significant decrease in the Form Factor (indicating a less circular morphology), and an increase in the Perimeter/Area ratio in cells treated with FGF2 compared to the other conditions (Fig. 4k, l). BGJ398 also induced a small, opposite, difference in these cell morphology parameters compared to DMSO Control treatment. No evidence of an effect of FGF2 on Ocy454 viability was observed (not shown) though a slight reduction in proliferation was noted following FGF2 treatment (Fig. S6b).These data are consistent with a role for FGF signaling in promoting dendritic outgrowth during osteocyte differentiation.

### *Fgfr* inactivation causes osteocyte apoptosis only in regions of new bone growth

Observations of TUNEL+ osteocytes, PDPN and OCN expression and loss of the LCN in FGFR1-CKO and FGFR1,2-DCKO mice were all restricted to osteocytes closer to the endosteum. To investigate if this was linked to new appositional bone growth occurring after inactivation of FGFR signaling, we performed a preliminary analysis to assess where bone formation was occurring under our experimental conditions. Control and FGFR1-CKO mice were exposed to tamoxifen beginning at 12 weeks of age for four weeks and injected with alizarin red 13 days and calcein green 3 days prior to harvest to mark the mineralizing front in any newly formed bone (Fig. S7a). Examination of longitudinal sections of the posterior tibial cortex from both Control and FGFR1-CKO mice showed clear alizarin-red labels within the bone near the endosteal surface, with parallel calcein green labels on the surface on the endosteal side (Fig. S7b). By contrast, on the periosteal surface only faint overlapping of alizarin and calcein labels were observed. This established that there was active appositional bone growth on the endosteal surface of Control and FGFR1-CKO tibias, but negligible bone apposition on the periosteal surface.

To analyze the relationship between newly formed bone and TUNEL+ osteocytes, 12-week-old Control mice harboring the DMP1-CreER; *ROSA^TDT^*alleles or floxed alleles of *Fgfr1* and *Fgfr2* (DFF), and FGFR1,2-DCKO mice were exposed to tamoxifen and administered a single dose of calcein green to mark the osteogenic growth front at the time of tamoxifen induction (Fig. 5a). Four weeks later, tibiae were harvested and analyzed for TUNEL and the extent of endosteal bone growth. Consistent with our initial experiments (Fig. 1), TUNEL+ osteocytes were seen only in FGFR1,2-DCKO cortical bone (Fig. 5b). Importantly, these TUNEL+ cells were restricted to the zone of newly formed bone on the endosteal side, whose outer boundary was delineated by calcein (green). To confirm this observation and to better distinguish between TUNEL+ osteocytes and the outer boundary of the newly formed bone, we repeated this study with FGFR1-CKO mice that did not harbor the *ROSA^TDT^* reporter allele and injected alizarin red instead of calcein green (see Fig. 5a). Like in Fig. 5b, we observed TUNEL+ osteocytes (green) only within the zone of newly formed endosteal bone whose boundary was marked by alizarin red (Fig. S8a). These data show that only osteocytes formed following induction of *Fgfr* inactivation are dying by apoptosis. This conclusion is consistent with the absence of any TUNEL+ cells, the absence of any PDPN/OCN-expressing osteocytes, and the intact LCN on the periosteal side, where there is negligible appositional bone growth (Figs. 2, 4).

**Figure 5.**
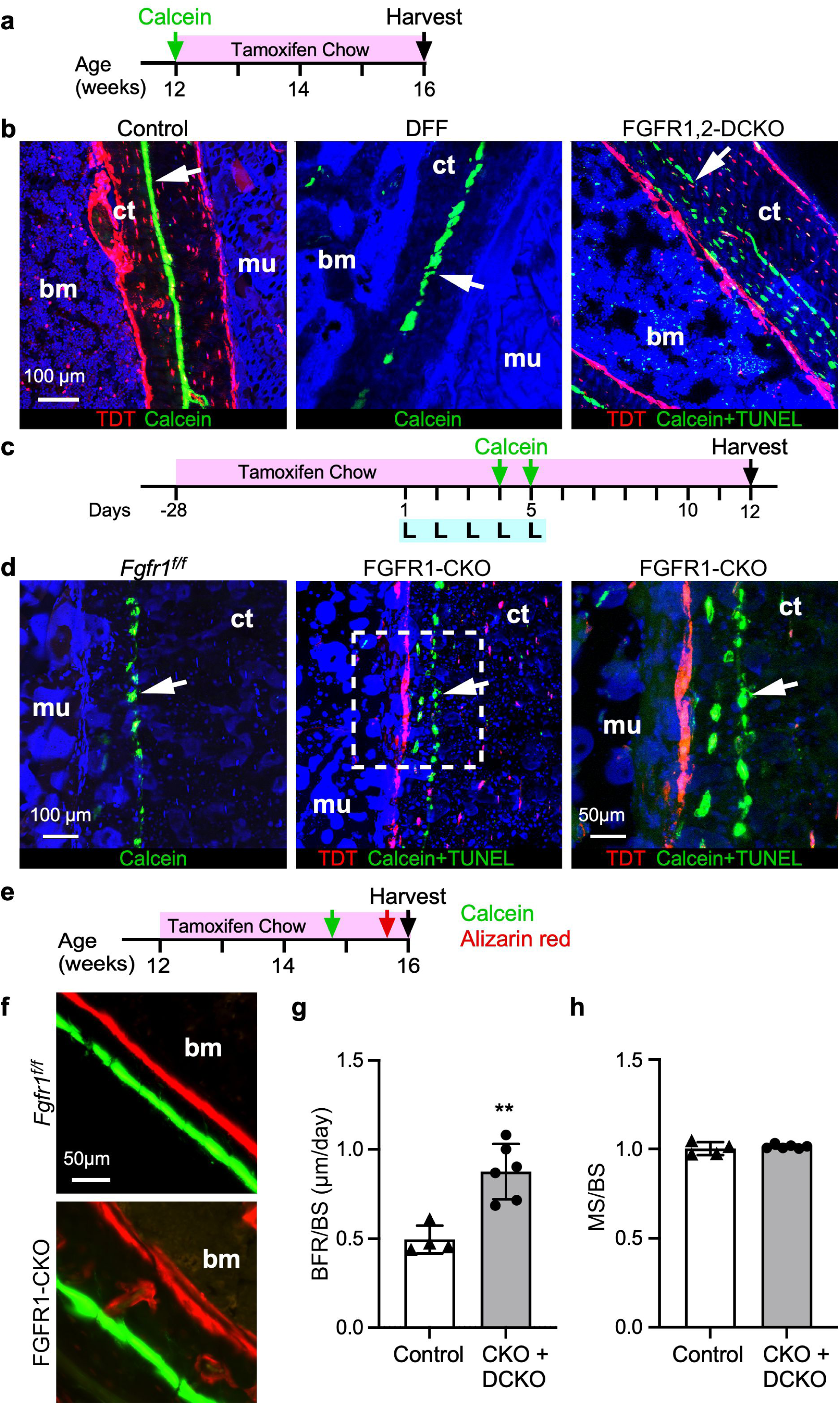
Osteocyte death is observed in newly formed cortical bone after *Fgfr* inactivation. **(a)** Experimental plan in which 12-week-old mice were exposed to tamoxifen from 12 to 16 weeks of age to induce DMP1-CreER and injected with calcein at 12 weeks of age to mark the boundary of the newly formed bone after the onset of *Fgfr1* inactivation. **(b)** The tibia of Dmp1-CreER; *ROSA^TDT^* (Control), *Fgfr1^f/f^; Fgfr2^f/f^* (DFF) and FGFR1,2-DCKO mice were assessed for apoptosis via TUNEL assay. TUNEL+ osteocytes were present only withing the boundaries of the newly formed bone from the endosteal side of FGFR1,2-DCKO mice. Arrow indicates calcein marking the outer boundary of newly formed bone from the endosteum (also see Fig. S8). *ROSA^TDT^* (red) marks the endosteum and periosteum in Control and FGFR1,2-DCKO mice (which harbor both DMP1-CreER and the *ROSA^TDT^* reporter alleles). Scale bar: 100 µm **(c)** Experimental plan to induce new bone formation by mechanical loading in the periosteal side of the tibia after *Fgfr1* inactivation. Mice were exposed to tamoxifen from 14 weeks of age until sacrifice. After 4 weeks of tamoxifen, the right tibia was loaded for five consecutive days and harvested 7 days after the end of loading. Calcein was injected on days 4 and 5 of loading to mark the newly formed bone. The non-loaded contralateral limb showed no periosteal appositional bone growth (not shown). **(d)** Images of TUNEL staining of *Fgfr1^f/f^* and FGFR1-CKO tibia after loading showing apoptosis in the newly formed osteocytes on the periosteal side of the posterior tibia in FGFR1-CKO mice. Arrow indicates the incorporation of calcein establishing the boundary of the newly formed bone. *ROSA^TDT^* (red) marks the periosteum with TUNEL+ osteocytes present in between these two boundaries. The dashed square outlines the region magnified on the right. Blue; DAPI-stained nuclei; bm, bone marrow; ct, cortical bone; mu, muscle. Scale bars: 50 µm (inset in (**d)**), 100 µm (**b, d**). **(e)** Experimental plan for dynamic histomorphometry on *Fgfr1^f/f^* and FGFR1-CKO mice induced with tamoxifen from 12 to 16 weeks of age and injected with calcein green 13 days and alizarin red 3 days, before sacrifice. **(f)** Representative images of calcein green and alizarin red incorporation in the endosteal side of the femur diaphysis of *Fgfr1^f/f^* and FGFR1-CKO mice. Scale bar: 50 µm. (**g, h**) Dynamic histomorphometry analysis was performed to calculate the bone formation rate/bone surface (BFR/BS) and mineralizing surface/bone surface (MS/BS) in Control (n=4), FGFR1-CKO (CKO, n=3) and FGFR1,2-DCKO (DCKO, n=3) mice. Data shown as the mean ± SD. Unpaired *t*-test of Control compared to combined CKO plus DCKO data, \*\**P*<0.01. Blue; DAPI-stained nuclei; bm, bone marrow; ct, cortical bone; mu, muscle.

To confirm that osteocyte apoptosis only occurs in regions of new appositional bone growth, we induced periosteal appositional bone growth by mechanical loading ^28^. In this model, mechanical loading of the tibia induces new periosteal bone growth on the posterior side of the tibia. Tamoxifen was administered to 14-week-old Control (*Fgfr1^f/f^*) and FGFR1-CKO mice for at least four weeks to induce *Fgfr* inactivation. Mice were then subjected to tibial loading (one session per day for 5 days) and given two doses of calcein on day 4 and 5 of loading to mark newly forming bone (Fig. 5c). Seven days later, tibiae were processed and analyzed for apoptosis. Control (*Fgfr1^f/f^*) and FGFR1-CKO mice showed new bone growth on the periosteal side of the posterior tibia, delineated by the calcein label embedded within the bone (Fig. 5d). This new growth was also accompanied by TUNEL+ osteocytes between the calcein (arrow) and periosteal (labeled red by TDT) boundaries in FGFR1-CKO mice, similar to that seen on the endosteal surface in the absence of any mechanical loading (Fig. 5b). Repeating this study with FGFR1,2-DCKO mice without the *ROSA^TDT^* reporter allele, and injection with alizarin red (instead of calcein green) on loading days 4 and 5 (see Fig. 5c) demonstrated the presence of TUNEL+ osteocytes (green) within newly formed bone (red) (Fig. S8b). Increased alizarin red incorporation suggests that FGFR1,2-DCKO could have a greater anabolic response to loading compared to FGFR1-CKO mice. These data confirm that only newly formed osteocytes, but not preexisting osteocytes, become TUNEL+ on *Fgfr* inactivation.

To further test whether *Fgfr1* loss can directly affect mature osteocytes, *Fgfr1* was inactivated with Sost-CreER in adult mice. Sost-CreER efficiently targets all mature osteocytes (even in the absence of tamoxifen) as reported previously ^29^ (Fig. S9a). Sost-CreER; *Fgfr1^f/f^* (SOST-FGFR1-CKO) mice show decreased expression of *Fgfr1* in flushed cortical bone (Fig. S9b) but showed no evidence of increased osteocyte death by apoptosis (Fig. S9c). Together with previous experiments, this suggests that only mature osteoblasts and newly forming osteocytes are dying in direct response to *Fgfr1* inactivation.

Finally, dynamic histomorphometry was used to quantify the effect of FGFR loss on bone growth. Tamoxifen was administered to 12-week-old mice and calcein green was injected 13 days and alizarin red injected 3 days before harvest (Fig. 5e, f). Both FGFR1-CKO and FGFR1,2-DCKO mice showed increased bone formation rate (BFR) rate compared to *Fgfr1^f/f^* and DFF control mice (Fig. 5g, h) but no difference in the mineralizing surface/bone surface ratio (Fig. S7c). These data show that loss of FGFR and osteocyte death is associated with increased endosteal bone formation.

### Gene expression analyses show similar responses to FGFR signaling in adult mouse cortical bone and in Ocy454 cells

To identify changes in gene expression in response to inactivation of *Fgfr1 in vivo*, femoral diaphyseal bone was harvested from 16-week-old Control (*Fgfr1^f/f^*) and FGFR1-CKO mice following 4 weeks exposure to Tamoxifen (Fig. 1a), flushed with PBS and subjected to bulk mRNA sequencing. Because of variability in tissue homogeneity and efficiency of gene inactivation, of the 6 Control (*Fgfr1^f/f^*) and 7 FGFR1-CKO samples sequenced, 4 Control samples with the highest *Fgfr1* expression were compared to 4 CKO samples with the lowest *Fgfr1* expression (Fig. 6a; Fig. S10a, b). With an adjusted *P*-value threshold set at <0.05 and a Log2FC threshold set at ≥0.5 or ≤-0.5, 1994 differentially expressed genes (DEG) were identified, with 1265 DEG lower and 729 DEG higher in FGFR1-CKO samples (Fig. 6b). Expression of several identified DEG that were decreased in FGFR1-CKO mice were validated by qRT-PCR on cortical bone RNA (Fig. S10c).

**Figure 6.**
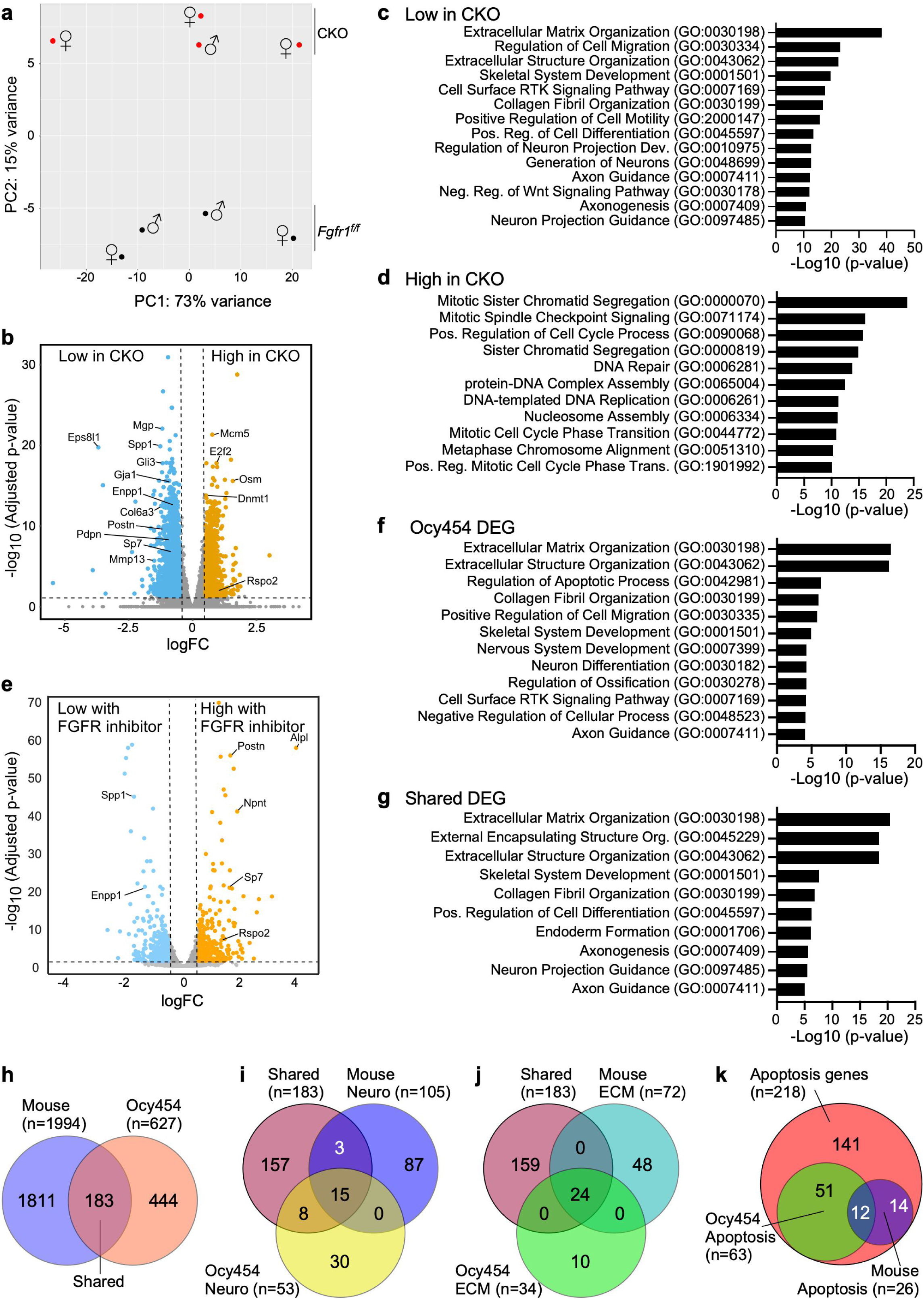
Differential mRNA analysis from *in vivo Fgfr* inactivation in mice and *in vitro* inhibition of FGFR signaling in Ocy454 cells. **(a)** Principal component analysis of differential mRNA expression harvested from the cortical hind limb bones from four Control (*Fgfr1^f/f^*) and four FGFR1-CKO mice after exposure to tamoxifen from 12-16 weeks of age (also see Fig. S10). **(b)** Volcano plot showing DEG in Control and FGFR1-CKO mice. Several representative DEG are indicated. Dashed lines are placed at *P*adj<0.05 and |log2FC| >0.5. (**c, d**) Pathway enrichment analysis of 1265 DEG that were downregulated in FGFR1-CKO mice (Log2FC ≥ 0.5, *P*adj <0.05) (**c**) and 729 DEG that were upregulated in FGFR1-CKO mice (Log2FC ≥ 0.5, *P*adj ≤0.05) (**d**). **(e)** Volcano plot of DEG in Ocy454 cells following treatment with FGF2 or BGJ398+FGF2, corresponding to panels (**i**) and (**j**) in Fig 4. Dashed lines are placed at *P*adj<0.05 and |log2FC| >0.5. **(f)** Pathway enrichment analysis of all DEG (|Log2FC| ≥ 0.5, *P*adj<0.05) identified in Ocy454 cells treated with FGF2 vs BGJ398+FGF2. **(g)** Pathway enrichment analysis of the 183 shared DEG from the *in vivo* mouse data and the *in vitro* Ocy454 cell culture data. **(h)** Venn diagram indicating the number of DEG shared between the *in vivo* mouse model and the *in vitro* Ocy454 model. (**I, j**) Venn diagram indicating the number of neuronal regulatory genes (**i**) and ECM genes (**j**) shared between the *in vivo* mouse model and the *in vitro* Ocy454 model. (**k**) Venn diagram showing unique and shared genes that regulate apoptosis.

Pathway enrichment analysis of DEG downregulated by *Fgfr* inactivation (Low in CKO) showed categories related to mature cortical bone such as Extracellular Matrix Organization, Skeletal System Development, and Collagen Fibril Organization (Fig. 6c; Fig. S10d). Interestingly, several categories included neuronal related terms such as Regulation of Neuron Projection Development, Generation of Neurons, and Axon Guidance (Fig. 6c; Fig. S10e). This is consistent with the known expression of neuronal genes in the formation of dendrites within the osteocyte LCN ^30–32^ and the observation of impaired osteocyte dendrite formation in FGFR1-CKO mice (Fig. 4a-e). In contrast, pathway enrichment analysis of DEG that were higher in FGFR1-CKO samples represented categories involved in cell cycle regulation and cell stress, such as Mitotic Sister Chromatid Segregation, DNA Repair and Mitotic Cell Cycle Phase Transition (Fig. 6d). This is consistent with increased expression of cell stress genes in mature FGFR1-CKO osteoblasts (Fig. S11a), and potential triggers for apoptosis in newly formed osteocytes; however, gene ontology terms related to apoptosis were not significantly enriched.

Bulk mRNA seq was also conducted on differentiated Ocy454 cells treated with FGF2, with or without the pan FGFR inhibitor, BGJ398 for 48 hr. With the adjusted *P*-value threshold set at <0.05 and a Log2FC threshold set at ≥0.5 or ≤-0.5, 627 DEG were identified, with 339 DEG lower in inhibitor-treated samples and 288 DEG higher in inhibitor treated samples (Fig. 6e). Pathway enrichment analysis of these DEG showed categories related to bone maturation such as Extracellular Matrix Organization, Collagen Fibril Organization and Skeletal System Development, as well as terms associated with neuron development such as Neuron Differentiation and Axon Guidance (Fig. 6f) similar to what was found in the mouse data. None of the cell stress related genes regulated in the mouse data (Fig. S11a) showed differential expression in Ocy454 cells (Fig. S11b), but the pathway Regulation of Apoptotic Process was enriched in Ocy454 cells (Fig. 6f).

We next compared all DEG from the mouse bone and Ocy454 data sets and found 183 shared DEG. Pathway enrichment analysis showed similar terms to those in the parental data sets including Extracellular Matrix Organization, Skeletal System Development and Axon Guidance (Fig. 6g, h). Of the 183 shared genes, there were 15 common neuronal regulatory genes, and 24 common ECM associated genes in both mouse and Ocy454 data (Fig. 6i, j; Fig. S11c, d). Prominent in the ECM list were several collagen genes and several *Adamts* (A disintegrin-like and metallopeptidase with thrombospondin type 1 motif) genes that are involved in modifying the ECM. The neuronal regulatory gene list includes the axon guidance genes *Slit3* and *Robo2* and several members of the *Sema3* gene family.

Because apoptosis is a predominant phenotype in FGFR1-CKO bone, we selectively evaluated expression of genes related to apoptosis. We curated a list of 218 apoptosis-related genes derived from GO terms and literature search and identified 26 DEG that include both pro-and anti-apoptotic genes in the mouse DEG list (Fig. 6K; Fig. S12a), reflecting the complex response to loss of FGFR signaling in different stages of the mature osteoblast lineage. A similar analysis of the Ocy454 data identified 63 pro- and anti-apoptosis DEG, with twelve apoptosis-related genes shared between the mouse and Ocy454 DEG sets. We visualized the expression of the 26 mouse DEG in the Ocy454 data set (Fig. S12b) and found that these genes tend to be less differentially regulated or counter-regulated between mouse and Ocy454 cells (Fig. S12c, d). Taken together, these data suggest that loss of signaling through FGF receptors does not directly induce apoptosis of osteoblast lineage cells. Instead, signaling through FGF receptors may be directly required for ECM remodeling and osteocyte projection formation, and loss of FGFR signaling causes retention of an osteoblast-like gene expression signature and apoptosis of bone-embedded cells.

### Structural alterations of the bone matrix on *Fgfr1* inactivation

Analysis of the RNA seq data revealed gene expression changes in apoptosis and ER stress pathways, which are in keeping with the observed osteocyte apoptosis when *Fgfr1* is inactivated in the osteoblast/osteocyte lineage (Fig. S10-12). Even though increased bone formation was observed by dynamic histomorphometry, RNA seq analysis revealed a significant downregulation of components of the ECM (Fig. S10d). To reconcile these observations, we investigated the ECM by histological and ultrastructural analyses. Toluidine blue staining and *von Kossa* staining revealed that the osteoid layer is greatly reduced in FGFR1-CKO and FGFR1,2-DCKO mice (Fig. 7). Toluidine blue staining identified a bluish-stained region next to the bone marrow, representing non-mineralized osteoid, which was greatly reduced in the FGFR1-CKO tibia and femur (Fig. 7a, b). Analysis of mineralization of the bone ECM with the *von Kossa* stain showed mineralized cortical bone (dark brown) in both Control and FGFR1,2-DCKO femur. The osteoid layer, which is not stained by *von Kossa* and lies adjacent to mineralized bone, was reduced in FGFR1,2-DCKO compared to Control femur (Fig. 7c). These data suggest abnormal mineralization of the bone ECM in the absence of FGFR signaling. Consistent with these histological stains, IF for COL1a1 in FGFR1-CKO and Lumican (a proteoglycan component of the bone matrix and marker that is expressed in differentiating osteoblasts^33^) in FGFR1,2-DCKO mice showed prominent staining in the endosteum of Control cortical bone and reduced staining in the corresponding CKO cortical bone (Fig. 7d, e). These observations collectively indicate that the osteoid is a short-lived structural component of the newly forming bone ECM and is consistent with altered mineralization of the osteoid in the absence of FGFR signaling. The is consistent with a downregulation of several genes associated with the bone mineralization process (Fig. S13a-c). IF for Osteomodulin (OMD), a leucine-rich proteoglycan that binds to hydroxyapatite crystals and collagen fibers ^34, 35^, showed a lamellar staining pattern in Control cortical bone, and a disorganized, woven bone pattern on the endosteal side of FGFR1,2-DCKO cortical bone (Fig. 7f), suggesting disorganization of bone ECM structure.

**Figure 7.**
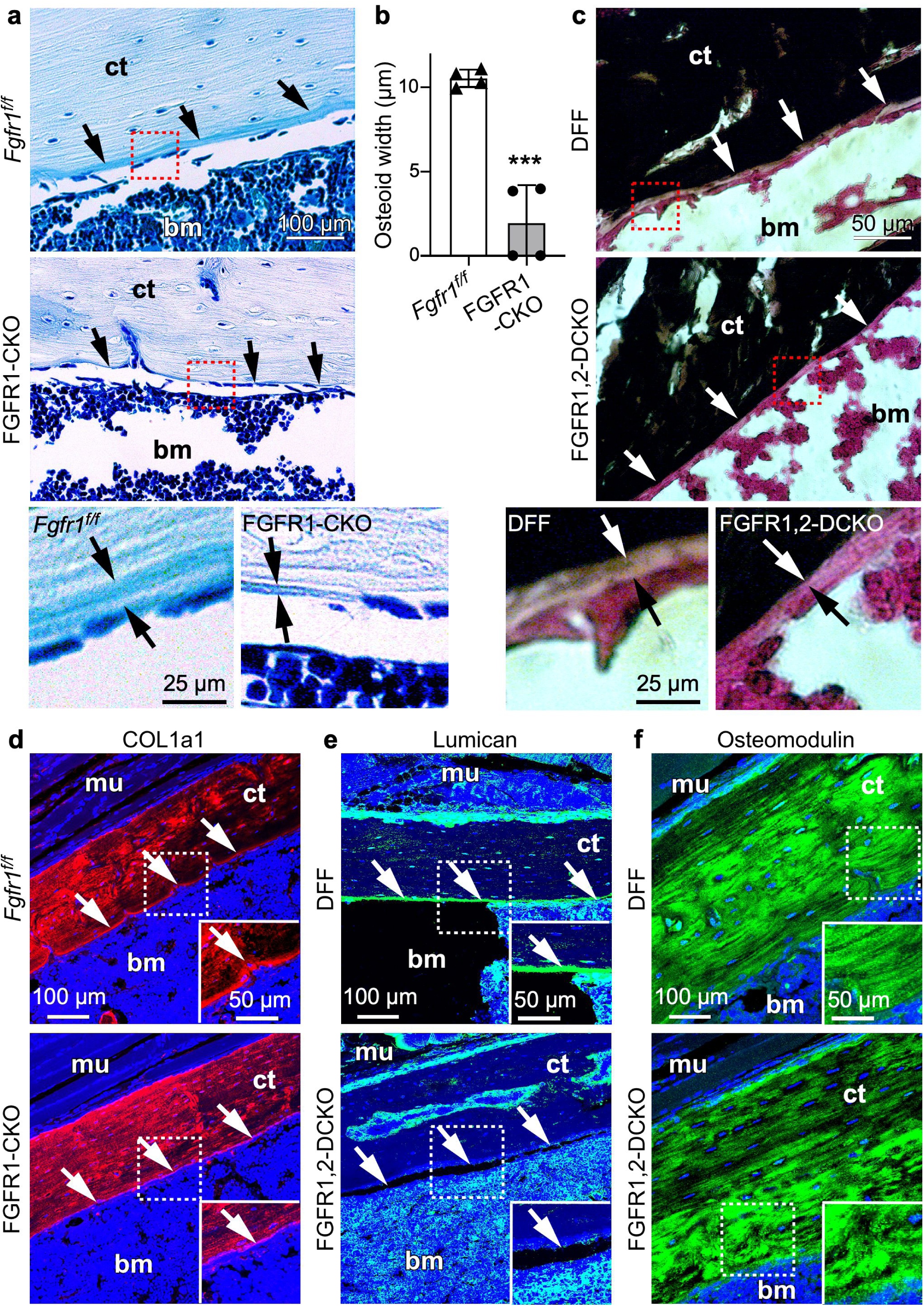
Abnormal osteoid development with *Fgfr* inactivation in the DMP1-lineage. Control, FGFR1-CKO and FGFR1,2-DCKO were treated with tamoxifen from 12 to 16 weeks of age and then sacrificed for histological analysis (see Fig. 1(b)). **(a)** Toluidine blue staining of the femur from *Fgfr1^f/f^* and FGFR1-CKO mice. The dashed boxes are magnified below. Arrows indicate the osteoid layer. **(b)** Bioquant analysis of endosteal osteoid layer thickness from *Fgfr1^f/f^* and FGFR1-CKO mice. *** *P*<0.001. **(c)** *von Kossa* staining of the femur from DFF and FGFR1,2-DCKO mice. The dashed boxes are magnified below. Arrows indicate the osteoid layer. (**d-f**) IF for COL1A1 (red) (**d**), Lumican (green) (**e**) and Osteomodulin (green) (**f**) in *Fgfr1^f/f^*, DFF Controls and FGFR1-CKO or FGFR1,2-DCKO mice. The dashed boxes are magnified in the insets. Arrows indicate the presence or absence of the osteoid in Control (*Fgfr1^f/f^* or DFF) and its reduction in FGFR1-CKO and FGFR1,2-DCKO mice. Blue; DAPI-stained nuclei; bm, bone marrow; ct, cortical bone. Bars: all panels, 100 µm; insets, 50 µm.

Scanning (SEM) and transmission (TEM) electron microscopy were used to examine the ultrastructure of FGFR1-CKO cortical bone. Disorganization of the cortical bone ECM on the endosteal side is seen as early as three weeks after tamoxifen induction of *Fgfr1* inactivation (Fig. 8) at which point TUNEL+ osteocytes are seen infrequently indicating that abnormal ECM is a major cause of osteocyte death. SEM of Control *Fgfr1^f/f^* tibia showed the presence of an organized lamellar banding with the collagen fibrils aligned parallel to the long axis of the bone, indicating normal organization of collagen fibrils in the ECM both in the osteoid region adjacent to the bone lining cells and deeper inside the bone on the endosteal side (Fig. 8a-c). In FGFR1-CKO mice, the collagen fibrils in the osteoid are arranged perpendicular to the axis of the bone with a reduction in lamellar banding indicating an absence of normal structural organization of the osteoid ECM (* in Fig. 8b), validating the abnormalities in osteoid staining seen in Figure 7. Deeper inside the bone, the ECM is disorganized with the collagen fibrils arranged randomly and a complete absence of lamellar banding (Fig. 8c). TEM analysis confirmed the disruption of the ECM in FGFR1-CKO mice with the appearance of irregular ridges that traversed the long axis of the bone, with a total lack of an organized homogenous pattern that is seen in Control ECM (Fig. 8 d, e). SEM and TEM analysis of Control and FGFR1-CKO tibia four weeks after tamoxifen induction demonstrated a similar lack of lamellar banding with irregular organization of collagen fibrils by SEM (Fig. S14a, b); TEM analysis demonstrated enhanced and increased irregular ridges surrounding dead osteocytes in FGFR1-CKO mice, in contrast to the homogenous pattern of Control mice (Fig. S14c, d).

**Figure 8.**
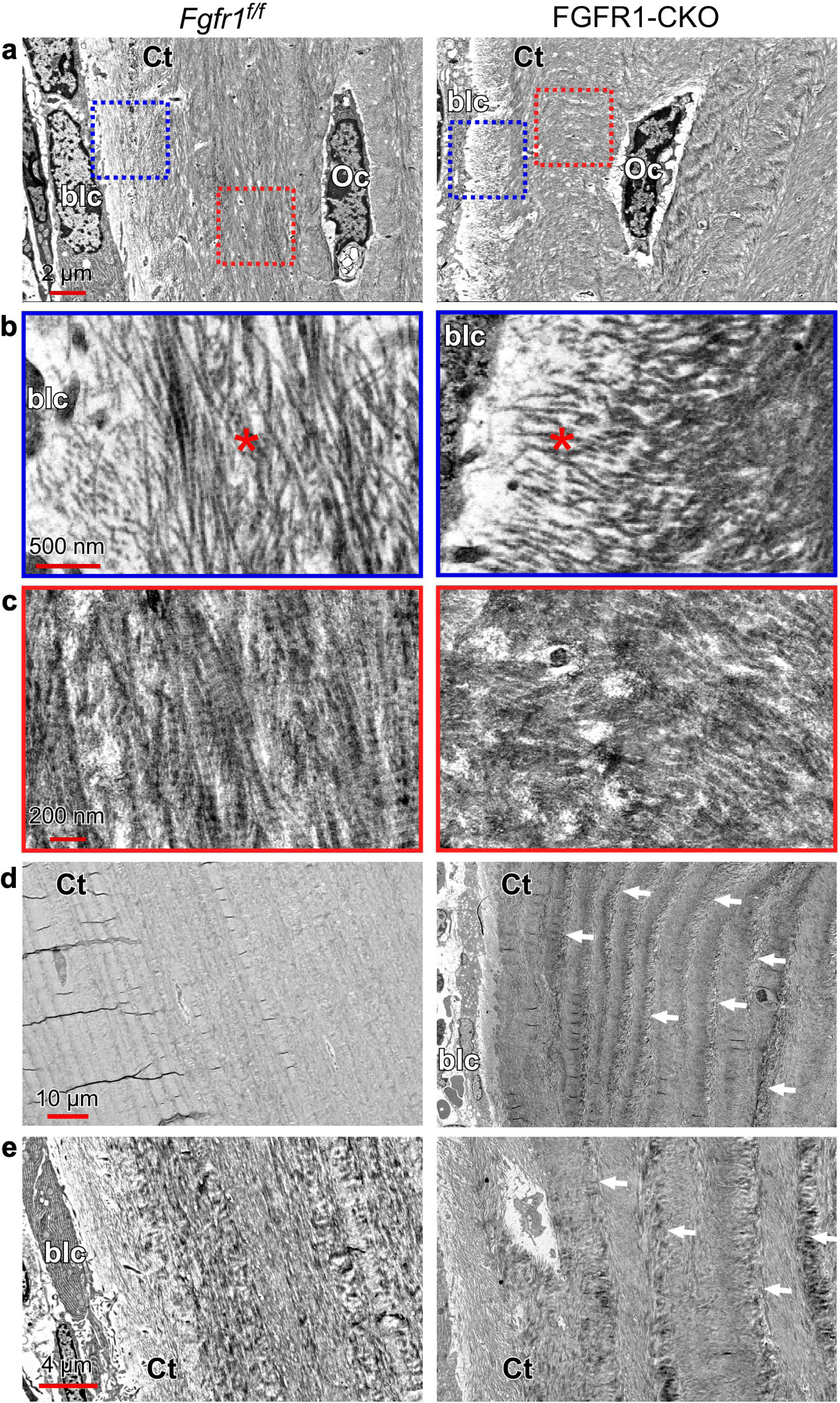
Abnormal ECM in regions of new bone formation following *Fgfr* inactivation in the DMP1 lineage after three weeks of tamoxifen induction. Mice were exposed to tamoxifen from 12-15 weeks of age and tibiae were harvested for ultrastructural analysis. (**a-c**) Scanning electron microscopy (SEM) images of *Fgfr1^f/f^*and FGFR1-CKO tibiae in regions of new endosteal bone formation under different magnifications. (**b)** Magnified image of the osteoid layer (*) next to bone lining cells (blc) (blue-dashed square) in (**a)**; (**c)** Magnified image of deeper structures within cortical bone (red-dashed square) in (**a)**. (**d, e**) Transmission electron microscopy (TEM) images of *Fgfr1^f/f^*and FGFR1-CKO tibiae in regions of new endosteal bone formation under different magnifications. Arrows indicate irregular ridges in the ECM of FGFR1-CKO tibia. blc, bone lining cell; Oc, osteocyte; ct, cortical bone. Scale bars: (**a**) 2 µm; (**b**) 500 nm; (**c**) 200 nm; (**d**) 10 µm; (**e**), 4 µm.

## Discussion

Osteocytes are long-lived cells that are embedded within mineralized bone matrix. The cell bodies of the osteocytes are housed in the lacunae with their dendritic processes projecting through the network of canaliculi, together forming the lacunocanalicular network (LCN), central to all osteocyte functions. In the adult, longitudinal bone growth stops, but appositional bone growth and bone remodeling continue, partly regulated by mechanical load. A primary osteocyte function throughout life is to regulate bone remodeling through signaling to osteoblasts to make new bone and to osteoclasts to resorb bone ^36^. The data presented here demonstrates that FGFR signaling in osteolineage cells modulates appositional bone growth in adult bone and has a vital role in osteocyte differentiation, survival and formation of the LCN complex.

Our findings demonstrate that lack of FGFR signaling disrupts the osteoblast to osteocyte transition and results in the ectopic expression of the late osteoblast markers, OCN and ALPL, and the early osteocyte transition marker E11/PDPN, in dying osteocytes in regions of appositional bone growth. E11/PDPN is essential for the formation of dendrites that form the LCN network ^13, 37, 38^. E11/PDPN expression in regions of dying osteocytes is suggestive of two different biological processes which are not necessarily mutually exclusive. First, impaired formation of the LCN could trigger a feedback mechanism that upregulates PDPN as a survival response by the dying osteocyte to loss of the LCN. The increased expression of stress response genes seen on *Fgfr* inactivation could be due to an increase in protein synthesis by such feedback mechanisms, Second, loss of FGFR signaling could lead to abnormal differentiation of late osteoblasts into osteocytes and retention of immature markers such as E11/PDPN and OCN. However, given the concurrent expression of PDPN, ALPL and OCN with the emergence of TUNEL+ osteocytes, it seems unlikely that expression of these proteins trigger osteocyte death. Regardless of the mechanism of cell death, dendritic outgrowth in Ocy454 cells in response to FGF is supportive of a direct role for FGFR signaling in osteocyte differentiation. The increase in E11/PDPN protein expression in TUNEL+ osteocytes is at odds with the observed decrease in *E11/Pdpn* mRNA seen in mRNA seq analysis of flushed cortical FGFR1-CKO bones. However, any cortical bone contains many different cell types, and several can express PDPN, particularly maturing osteoblasts and bone lining cells. It is possible that in Control bone *Pdpn* mRNA is expressed at higher levels in osteoblasts but not in osteocytes; in FGFR1-CKO tissue *Pdpn* mRNA could be lower in the osteoblasts, but PDPN protein epitopes could be retained in the lacunae of dying osteocytes.

A key phenotypic manifestation of the loss of FGFR signaling in adult bone is the specific localization of TUNEL+ osteocytes in the endosteal region, which is the site of baseline bone apposition under normal loading conditions. This pattern of osteocyte death in cortical bone strongly co-localizes with the expression of PDPN, ALPL and OCN in regions where there is loss of the LCN network. Delineation of the region of new appositional bone growth with calcein or alizarin red labeling of the mineralization front shows that at this stage all TUNEL+ and PDPN-, ALPL- and OCN-expressing osteocytes lie within newly formed bone matrix, indicating that the underlying mechanism of osteocyte death in the absence of FGFR signaling is linked to bone matrix formation. This correlation is further strengthened by the observation that TUNEL+ osteocytes can be seen in the periosteal region when mechanical loading induces new bone formation on this surface.

In the absence of FGFR signaling several marked perturbations in the bone ECM were observed that suggest that the newly formed bone matrix is not conducive to supporting osteocyte viability. The osteoid layer in newly forming bone was significantly reduced in the absence of FGFR signaling indicating reduced synthesis and/or altered mineralization of the newly deposited osteoid matrix. During the osteoblast to osteocyte transition, the newly differentiating osteocytes normally embed in the osteoid and it is here that the cell first elaborates dendritic projections that eventually form the LCN ^10^. Despite the reduced thickness of the osteoid layer, dynamic histomorphometry showed increased bone formation within 4 weeks when *Fgfr1*was inactivated in 12-week-old mice, which may contribute to the dysregulation of mineralization in the newly formed bone. The long-term effects of loss of FGFR1 in adult differentiating osteoblasts will be an interesting area for future investigation.

The osteoid layer in FGFR-deficient bone is also remarkably reduced in bone ECM components such as COL1a1 and Lumican. Perturbation of bone ECM is also seen in the disorganized deposition of OMD (Osteomodulin/Osteoadherin) close to the endosteum. OMD is important for the deposition of collagen fibrils and mice lacking OMD have a reduced endosteal and periosteal diameter in transverse sections ^35, 39^. Consistent with this disorganized staining pattern, ultrastructural analysis is also indicative of a poorly formed bone ECM in the absence of FGFR signaling. Thin section SEM and TEM show distorted, non-uniform bone ECM in the osteoid region and in deeper structures of the endosteal bone, suggestive of collagen fibrils that are either non-aligned, or poorly crosslinked. This is especially striking as these ECM distortions are widespread under conditions where TUNEL+ osteocytes are only beginning to appear. These observations are suggestive of ECM distortions driving osteocyte cell death. Taken together, these observations suggest that an osteoid layer is supportive or even necessary for osteocyte dendrite formation and the abnormal mineralization of osteoid, the deficiency of bone ECM components, and the accompanying ECM distortions would prevent dendrite formation and the formation of a functional LCN network resulting in impaired osteocyte viability within the newly formed bone ^40, 41^.

Our use of the DMP1-CreER driver to inactivate *Fgfr1* signaling targets both the transitioning osteoblast and the newly formed osteocyte. Using a Sost-CreER driver, which preferentially targets mature osteocytes, to inactivate *Fgfr1* does not generate a comparable phenotype indicating that the requirement for FGFR signaling lies either in the transitioning osteoblast or the nascent osteocyte. It is well established that active osteoblasts secrete collagenous bone ECM, but on differentiation, this ability is greatly reduced ^5, 36, 42^. The extent to which osteocytes produce bone ECM is not known; however, there is evidence *in vivo* that osteocytes can remodel the mineral content of the surrounding ECM under some conditions such as during lactation ^43^, produce at least some of the collagen ECM surrounding their lacunae, and influence collagen alignment in the existing collagen matrix ^24^. Currently, a definitive pathway delineating bone ECM formation and changes directly regulated by osteocytes remains to be elucidated, but our studies indicate that FGFR1 signaling is a vital part of this pathway regulating bone homeostasis.

## Materials and Methods

### Mice

All mice (*Mus musculus*) were housed in a pathogen-free barrier facility and handled in accordance with standard user protocols, animal welfare regulations, and the NIH Guide for the Care and Use of Laboratory Animals. All studies were performed under a protocol approved by the Institutional Animal Care and Use Committee at Washington University in Saint Louis.

*Fgfr1^f/f^* ^44^, *Fgfr2^f/f^* ^45^, DMP1-CreER ^44^, Sost-CreER ^29^ and the *ROSA26-tdTomato Ai9 allele (ROSA^TDT^*) ^46^ mice have been previously described. Mice were of mixed sexes and maintained on a mixed C57BL/6J × 129X1/SvJ genetic background. Homozygous floxed alleles of *Fgfr1* and *Fgfr2* were maintained as single (*Fgfr1^f/f^*, *Fgfr2^+/+^*) or double (*Fgfr1^f/f^*, *Fgfr2^f/f^*) floxed mice and in some cases additionally homozygous for the *ROSA^TDT^* allele. DMP1-CreER; *Fgfr1^f/f^*mice were maintained by crossing DMP1-CreER; *Fgfr1^f/f^* males to *Fgfr1^f/f^* or *Fgfr1^f/f^*; *ROSA^TDT/TDT^* females, resulting in a 50% yield of experimental and *Fgfr1^f/f^* Control mice. A similar mating scheme was used to generate DMP1-CreER; *Fgfr2^f/f^*mice and DMP1-CreER; *Fgfr1^f/f^*; *Fgfr2^f/f^* mice. Cre activation in DMP1-CreER; *Fgfr1^f/f^* (FGFR1-CKO), DMP1-CreER; *Fgfr2^f/f^*(FGFR2-CKO) or DMP1-CreER; *Fgfr1^f/f^*; *Fgfr2^f/f^* (FGFR1,2-DCKO) mice was induced for specific periods of time (1, 2, 3 or 4 weeks) by replacement of regular chow with tamoxifen chow (400mg/kg, Envigo Teklad, Madison, WI, USA; TD.130860); *Fgfr1^f/f^*, *Fgfr2^f/f^*, or *Fgfr1^f/f^*;*Fgfr2^f/f^*(DFF) mice were also identically treated as Controls. Sost-CreER; *Fgfr1^f/f^*; *ROSA^TDT^* mice were similarly created and the Sost-CreER similarly activated by tamoxifen and analyzed. In our experience, Sost-CreER appears to be constitutively active in osteocytes in the absence of tamoxifen, as previously observed ^29^.

### Histology, immunofluorescence staining, and imaging

Long bones (intact femurs and tibias) were isolated, fixed in 10% neutral buffered formalin for 24 hr at room temperature. After rinsing with PBS, bones were decalcified in 14% EDTA for 2 weeks and processed for paraffin embedding. Formalin-fixed paraffin-embedded (FFPE) sections were stained with Safranin O, Fast Green, Toluidine blue, and Hematoxylin & Eosin for morphological analysis or processed for silver staining ^47^ for analysis of the lacunocanalicular network (LCN) ^47, 48^. For observation of incorporated calcein green/alizarin red, tissue sections were fixed with 4% paraformaldehyde and further processed for 30% sucrose infiltration without decalcification, followed by embedding in OCT (Tissue-TekR). Mineralized tissues were also used for *von Kossa* staining.

For immunofluorescence (IF) on FFPE sections, antigen retrieval was done by overnight treatment of rehydrated FFPE sections with 10 mM citrate buffer (pH 6.0) at 60°C followed by gradual cooling to room temperature. After treating the sections with 0.3% Triton-X-100/PBS for 10 min. at room temperature, they were incubated with 1:100 dilution of the primary antibody at 4°C overnight. The following primary antibodies were used: Podoplanin (Abcam 11936), Osterix (Abcam 227820), Osteocalcin (Abcam 93876), mSost (RD Systems, AF1589), Alkaline phosphatase (Takara M190, Novus Biologicals AF2910), type I Collagen (Abcam 21286), Lumican (Abcam 168348). IF analysis of TDTomato (Kerafast 16D7 EST203), type I Collagen (Abcam 21286) and Lumican (Abcam 168348) expression on FFPE sections were done on rehydrated sections treated with proteinase K (10 ng/µl) at 37°C for 20 min. for antigen retrieval. The secondary antibodies were Alexa Fluor 488-goat anti-rabbit, Alexa Fluor 594-goat anti-rabbit, Alexa Fluor 594-goat anti-rat, Alexa Fluor 594-, or 488-goat anti-Syrian hamster (Life Technologies, Carlsbad, USA), at 1:200-1:250 dilution followed by counterstaining of the nuclei with DAPI. Analyses for apoptosis were performed on FFPE sections using the DeadEnd^TM^ Fluorometric TUNEL System staining kit (Promega, G3250) and manufacturer’s instructions provided. Images of stained bone sections were captured at high resolution using a Nikon A1/A1R confocal microscope. Z-stacks of images were taken of a minimum of 10 µm thickness followed by processing for maximum intensity projection. Canvas^TM^ X Draw was used for further image processing.

### Ultrastructural Analysis of bone by electron microscopy

Tibia from Control and FGFR1-CKO and FGFR1,2-DCKO mice were fixed overnight at 4°C in 2% paraformaldehyde/2.5% glutaraldehyde solution in 1X Hanks Buffered Salt Solution (HBSS) pH 7.4 and then decalcified in 14% EDTA for 2 weeks. Samples were then rinsed in HBSS buffer three times for 10 min each and subjected to a secondary fixation for one hour in 2% osmium tetroxide/1.5% potassium ferrocyanide in 1X HBSS. Following this, samples were rinsed in ultrapure water 3 times for 10 min. each and stained overnight in an aqueous solution of 1% uranyl acetate at 4°C. After staining was complete, samples were washed in ultrapure water 3 times for 10 min each, dehydrated in a graded acetone series (30%, 50%, 70%, 90%, 100%), four times in each solution for 10 min each. Next, samples were infiltrated with Spurr’s resin (30%, 50%, 70%, 100%) for 6 hr each, followed by 100% overnight and fresh 100% resin for 3 hr followed by fresh resin. Samples in 100% resin were heated in the microwave2 times for 5 min each at 30°C, (100% power, 350 Watts under vacuum (20 in/hg), (Pelco BioWave Pro, Redding, CA). Samples were then cured in an oven at 60°C for 48-72 hr. Post-curing, resin-embedded samples were oriented and mounted for longitudinal sectioning. 70 nm thick sections were cut from sample blocks and picked up on grids, post-stained with uranyl acetate and Reynolds lead ^49^, and then imaged on a JEOL JEM-1400 Plus (Tokyo, Japan) transmission electron microscope (TEM) operated at 120 KeV. Micrographs were made using an AMT (Nanosprint15-MkII) sCMOS camera. For scanning electron microscopy (SEM) imaging, 100 nm sections were cut and picked up on 10 x 10 mm silicon chips (Ted Pella Inc. Redding, Ca), post-stained with uranyl acetate and Reynolds lead and imaged in a FE-SEM (Zeiss Merlin, Oberkochen, Germany) using the ATLAS (Fibics Inc., Ottawa, Canada) scan engine to tile regions of interest. The FE-SEM was operated at 8 KeV and 300 pA using the solid-state backscatter detector.

### *In vivo* mechanical loading

To stimulate periosteal bone formation, *in vivo* mechanical loading (axial tibial compression) was applied as per an established protocol with slight adjustments ^28^. 14-week-old *Fgfr1^f/f^*, *Fgfr1^f/f^; Fgfr2^f/f^* (DFF), FGFR1-CKO and FGFR1,2-DCKO mice were exposed to tamoxifen chow for 4 weeks to induce *Fgfr* inactivation. Following this, under anesthesia (1-3% isoflurane), 11-N of peak force was applied via force-Controlled cyclic loading (Electropulse 1000; Instron) to the right hindlimb; left limbs served as contralateral non-loaded Controls. Sixty cycles of a 4-Hz sinusoidal waveform were applied daily for 5 days. In a preliminary cohort of Control and CKO mice, we confirmed that this loading protocol increased measures of bone formation on the periosteal surface determined by classic dynamic histomorphometric analysis (data not shown).

Buprenorphine (0.1 mg/kg) was injected subcutaneously after each loading bout for pain management. Following loading, mice were allowed free cage activity. Intraperitoneal injections of calcein green (10 mg/kg: Millipore Sigma) were administered on days 4 and 5 after loading. Mice were allowed normal activity for six days after the last loading and euthanized on the seventh day by CO_2_ asphyxiation. Limbs were harvested, formalin-fixed, infiltrated with 30% sucrose, frozen-embedded in OCT and sectioned longitudinally for analysis of TUNEL+ osteocytes in the newly incorporated periosteal bone on the posterior side of the tibia, the region where maximal loading-induced bone formation is observed in this loading model.

### Dynamic histomorphometry

To analyze the dynamic relationship between bone growth and osteocyte apoptosis, 12-week-old mice were injected intraperitoneally with calcein green (10 mg/kg, SIGMA, St. Louis, MO, USA) or alizarin red (30 mg/kg, SIGMA, St. Louis) on the day of DMP1-Cre activation with tamoxifen chow. Mice were harvested 4 weeks later, the limbs were fixed in 4% PFA at room temperature for 24 hr and processed for frozen sections after 30% sucrose infiltration (see above) using tape transfer ^50^ without decalcification. TUNEL analysis was performed after air-drying of frozen sections at room temperature for 2 hr using the DeadEnd^TM^ Fluorometric TUNEL System staining kit (Promega Inc.). The presence of TUNEL+ osteocytes in relationship to the osteogenesis front (visible by calcein/alizarin red incorporation) were observed by confocal fluorescence microscopy.

To measure mineral apposition (MAR) and bone formation (BFR) rates, a separate cohort of 12-week-old mice were exposed to tamoxifen chow for four weeks. Mice were injected with calcein intraperitoneally ten days before harvest and alizarin red three days before harvest. Hindlimbs were harvested, fixed with 4% PFA/PBS at room temperature for 24 hr and then processed as described above. Sections were scanned by fluorescence microscopy to capture calcein and alizarin red incorporation into bone and MAR and BFR were calculated using Bioquant Osteo software (www.bioquant.com).

### Treatments for *in vitro* experiments with Ocy454 cell line

The pan-FGFR1/2/3 inhibitor, BGJ398 (Selleck Chemical, S2183), was reconstituted in DMSO at 1 mM and diluted 1:100 in media to create a working solution of 10 µM. DMSO was diluted 1:100 in media to create a working solution of vehicle. Working solutions were diluted 1:50 into media for a final treatment dose of 200 nM BGJ398. FGF2 (Proteintech Group Inc., 50-197-3960) was reconstituted in PBS + 0.1% BSA at 100 µg/mL, then diluted into media for final treatment doses of 2 ng/mL or 5 ng/mL.

### Ocy454 cell culture

Ocy454 cells proliferate at 33°C under control of a temperature-sensitive large T antigen, and they stop dividing and begin to differentiate to an osteocyte phenotype at 37°C ^25, 26^. Alpha-MEM supplemented with 10% FBS (Gibco, A5256701) and 1% antibiotic-antimycotic (Gibco, 15240062) media was used for all experiments. Ocy454 cells were expanded at 33°C, 5% CO_2_, in a humidified incubator. Ocy454 cells were then plated into glass chamber slides (Corning Life Sciences, 354118) in complete media at a density of 2000 cells per well.. After 24 hr, treatment media was applied for 72 hr for each of four conditions: DMSO, DMSO + FGF2 (5 ng/mL), BGJ398, BGJ398 + FGF2 (5 ng/mL). At the end of the treatment period, cells were fixed in 4% PFA, permeabilized in 0.05% saponin, and stained with phalloidin (Abcam Cytopainter, ab112125) and DAPI to visualize the F-actin cytoskeleton. Cells were imaged on a Nikon Eclipse microscope with a 20x objective. Changes in Ocy454 cell morphology were quantified with a custom pipeline using CellProfiler software ^27, 51, 52^. Briefly, nuclei were identified from single channel DAPI images (primary objects), and cell bodies corresponding to each nucleus were identified from single channel phalloidin images (secondary objects). The experiment was repeated at least 3 times with representative images shown.

### mRNA sequencing of Ocy454 cells

Ocy454 cells were grown to confluence and then allowed to differentiate at 37°C. After 5 days, cells were pretreated for one hour with the pan-FGFR1/2/3 inhibitor BGJ398 or with DMSO vehicle. Treatment media was then applied for 48 hr for each of four conditions in triplicate: DMSO, DMSO + FGF2 (2 ng/mL), BGJ398, BGJ398 + FGF2 (2 ng/mL). On differentiation day 7, cells were harvested in guanidine-isothiocyanate lysis buffer containing 1% 2-mercaptoethanol. Lysates were passed through QIAshredder columns (Qiagen) RNA isolated and purified with PureLink RNA columns (PureLink RNA Mini Kit, Invitrogen) according to manufacturer’s instructions. 2 µg of purified RNA was submitted to Novogene for RNA library preparation with poly A enrichment and mRNA sequencing (NovaSeq, PE150). Reads were pseudo-aligned to the mouse transcriptome (Ensembl GRCm39 release 110) using Salmon ^53, 54^. Differential expression analysis of BGJ398+FGF2 samples compared to DMSO+FGF2 samples was performed with DESeq2 ^54^ with thresholding to include only genes with at least 10 counts in all samples. RNA sequencing data is available from the NCBI Gene Expression Omnibus, accession number GSE315887.

### mRNA sequencing of adult mouse cortical bone

DMP1-CreER*; Fgfr1^f/f^* (FGFR1-CKO) and Control *Fgfr1^f/f^*mice were induced with Tamoxifen chow from 12 to 16 weeks of age, and the femora and tibia were isolated at 16 weeks. Each diaphysis was dissected, the marrow was flushed with PBS with a 26G needle, and the remaining cortical bone was snap frozen in liquid nitrogen and stored at −80°C until further processing. Frozen tissues were pulverized in a liquid nitrogen-cooled stainless-steel flask with a ball bearing in a Micro Dismembrator (Sartorius Inc.) at 2000 rpm for 20 40 sec. RNA was stabilized with Trizol Reagent (Life Technologies Corporation, USA) and total RNA was isolated with the RNeasy Mini Kit (Qiagen Inc., Germantown, MD) according to the manufacturer’s instructions.

Six Control samples and seven FGFR1-CKO samples with RIN scores between 7.1 and 8.4 were sequenced at the Washington University Genome Technology Access Center (GTAC). Total RNA integrity was determined using Agilent 4200 TapeStation. Library preparation was performed with 200-500 ng of total RNA. Ribosomal RNA was removed by RNase-H using RiboErase kit (Roche). mRNA was then fragmented in reverse transcriptase buffer and heated to 94°C for 8 min. mRNA was reverse transcribed to yield cDNA using SuperScript III RT enzyme (Life Technologies) per manufacturer’s instructions and random hexamers. A second strand reaction was performed to yield ds-cDNA. cDNA was blunt ended, an A base added to the 3’ ends, followed by ligation of Illumina sequencing adapters. Ligated fragments were then amplified for 15 cycles using primers incorporating unique dual index tags. Fragments were sequenced on an Illumina NovaSeq 6000 using paired end reads extending 150 bases. Reads were processed using an in-house pipeline and open-source R packages. Briefly, raw reads were first trimmed using cutadapt to remove low quality bases and reads. Trimmed reads were then aligned to the mouse genome mm10 with GENCODE annotation vM25 using STAR (v2.7.9a) with default parameters. Transcript quantification was performed using featureCounts from the subread package (v2.0.1). Further quality control assessments were made using RSeQC and RSEM, and batch correction was performed using edgeR, EDASeq, and RUVSeq. RNA sequencing data will be deposited in the NCBI Gene Expression Omnibus (accession number GSE316171).

Principle component analysis and differential expression analysis for Control and FGFR1-CKO samples were determined using DESeq2 in negative binomial mode using batch-corrected transcripts from featureCounts (> 2-fold expression change, > 1 count per million (CPM), Benjamini corrected P < 0.05, and thresholding to include only genes with at least an average of 5 counts in all samples). Pairwise comparisons were made between Control vs FGFR1-CKO to determine differentially expressed genes (DEG). Initial analysis showed variability in the expression level of *Fgfr1* in Control and FGFR1-CKO samples, likely due to variability in marrow flushing, soft tissue adherence to the periosteal surface, and potential variability in knockout efficiency. To select samples for further analysis the read counts per exon for each of the 18 exons in *Fgfr1* were calculated for each sample. The average read counts for exons 2-17 were then compared across samples using a one-way ANOVA analysis with Tukey’s multiple comparisons test. Of the 78 pairwise comparisons between samples, 25 showed a statistical significance >0.05. Each sample pair was scored, with the higher read count sample given a +1 and the lower read count sample given a -1. Each sample was then ranked based on the sum of the significance score. This ranking correlated with ranking based on average mean read count for *Fgfr1* exons per sample. The top four ranking Control samples and the bottom four ranking CKO samples were used for further analysis. Gene ontology analysis was performed using Enrichr. Heatmaps were generated in R using Pheatmap (https://github.com/raivokolde/pheatmap).

### Comparison of RNA seq data sets

Venn diagrams were generated using the BioVenn web application ^55^. The list of 218 apoptosis related genes was derived from apoptosis gene ontology terms, AI generated list of apoptosis genes, and Ocy454 DEG found in the Regulation of Apoptotic Process (GO:0042981) gene ontology term.

## Acknowledgments

We thank Helen McNeil and her laboratory for the generous use of their Confocal Microscope (Nikon A1); Crystal Idleburg and Samantha Coleman of the Musculoskeletal Research Center (MRC) histology core and Marina Platik and Kathleen Lutker of the Washington University Histology Service and Innovation Center (WHISC) for help with sample preparation, sectioning and staining. We thank John Wulf II and Gregory Strout of the Washington University Center for Cellular Imaging (WUCCI) for their help with SEM and TEM. We thank Lynda Bonewald for sharing the Sost-CreER mouse line.

## Author contributions

D.P., C.S., C.M.M., M.N.W., M.J.S., and D.M.O. conceptualized and designed the study; D.P., C.S., C.M.M., M.N.W., M.J.S., and D.M.O. developed methodology; D.P., C.S., C.W., C.M.M. and I.A. performed the experiments; All authors analyzed the data; D.P. and D.M.O. drafted the manuscript; all authors reviewed and edited the manuscript; D.P., M.N.W., M.J.S. and D.M.O. acquired funding.

**Supplementary information accompanies the manuscript on the Bone Research website** http://www.nature.com/boneres.

## References

1. Manolagas, S.C. Birth and death of bone cells: basic regulatory mechanisms and implications for the pathogenesis and treatment of osteoporosis. Endocr Rev 21, 115–137 (2000).

2. Robling, A.G. & Bonewald, L.F. The Osteocyte: New Insights. Annu Rev Physiol 82, 485–506 (2020).

3. Wang, J.S. et al. Control of osteocyte dendrite formation by Sp7 and its target gene osteocrin. Nat Commun 12, 6271 (2021).

4. Cui, J., Shibata, Y., Zhu, T., Zhou, J. & Zhang, J. Osteocytes in bone aging: Advances, challenges, and future perspectives. Ageing Res Rev 77, 101608 (2022).

5. Schaffler, M.B., Cheung, W.Y., Majeska, R. & Kennedy, O. Osteocytes: master orchestrators of bone. Calcif Tissue Int 94, 5–24 (2014).

6. Xiong, J. et al. Matrix-embedded cells control osteoclast formation. Nat Med 17, 1235–1241 (2011).

7. Nakashima, T. et al. Evidence for osteocyte regulation of bone homeostasis through RANKL expression. Nat Med 17, 1231–1234 (2011).

8. Murali, S.K. et al. FGF23 Regulates Bone Mineralization in a 1,25(OH)2 D3 and Klotho-Independent Manner. J Bone Miner Res 31, 129–142 (2016).

9. Huang, X., Jiang, Y. & Xia, W. FGF23 and Phosphate Wasting Disorders. Bone Res 1, 120–132 (2013).

10. Dallas, S.L., Prideaux, M. & Bonewald, L.F. The osteocyte: an endocrine cell … and more. Endocr Rev 34, 658–690 (2013).

11. Dallas, S.L. & Veno, P.A. Live imaging of bone cell and organ cultures. Methods Mol Biol 816, 425–457 (2012).

12. Wetterwald, A. et al. Characterization and cloning of the E11 antigen, a marker expressed by rat osteoblasts and osteocytes. Bone 18, 125–132 (1996).

13. Zhang, K. et al. E11/gp38 selective expression in osteocytes: regulation by mechanical strain and role in dendrite elongation. Mol Cell Biol 26, 4539–4552 (2006).

14. Ornitz, D. FGF signaling in the developing endochondral skeleton. Cytokine Growth Factor Rev 16, 205–213 (2005).

15. Ornitz, D.M. & Itoh, N. Fibroblast growth factors. Genome Biology 2, REVIEWS3005 (2001).

16. Ornitz, D.M. & Itoh, N. New developments in the biology of fibroblast growth factors. WIREs Mech Dis 14, e1549 (2022).

17. Ornitz, D.M. & Marie, P.J. Fibroblast growth factor signaling in skeletal development and disease. Genes Dev 29, 1463–1486 (2015).

18. Jacob, A.L., Smith, C., Partanen, J. & Ornitz, D.M. Fibroblast growth factor receptor 1 signaling in the osteo-chondrogenic cell lineage regulates sequential steps of osteoblast maturation. Dev Biol 296, 315–328 (2006).

19. Xiao, Z. et al. Osteocyte-specific deletion of Fgfr1 suppresses FGF23. PLoS One 9, e104154 (2014).

20. Nookaew, I. et al. Refining the identity of mesenchymal cell types associated with murine periosteal and endosteal bone. J Biol Chem 300, 107158 (2024).

21. McKenzie, J. et al. Osteocyte Death and Bone Overgrowth in Mice Lacking Fibroblast Growth Factor Receptors 1 and 2 in Mature Osteoblasts and Osteocytes. J Bone Miner Res 34, 1660–1675 (2019).

22. van Bezooijen, R.L. et al. Sclerostin is an osteocyte-expressed negative regulator of bone formation, but not a classical BMP antagonist. J Exp Med 199, 805–814 (2004).

23. Bellows, C.G., Reimers, S.M. & Heersche, J.N. Expression of mRNAs for type-I collagen, bone sialoprotein, osteocalcin, and osteopontin at different stages of osteoblastic differentiation and their regulation by 1,25 dihydroxyvitamin D3. Cell Tissue Res 297, 249–259 (1999).

24. Shiflett, L.A. et al. Collagen Dynamics During the Process of Osteocyte Embedding and Mineralization. Front Cell Dev Biol 7, 178 (2019).

25. Spatz, J.M. et al. The Wnt Inhibitor Sclerostin Is Up-regulated by Mechanical Unloading in Osteocytes in Vitro. J Biol Chem 290, 16744–16758 (2015).

26. Wein, M.N. et al. HDAC5 controls MEF2C-driven sclerostin expression in osteocytes. J Bone Miner Res 30, 400–411 (2015).

27. Russ, J.C., F. Brent Neal The image processing handbook, Edn. 7th. (CRC/Taylor and Francis, Boca Raton; 2017).

28. Harris, T.L. & Silva, M.J. Dmp1 Lineage Cells Contribute Significantly to Periosteal Lamellar Bone Formation Induced by Mechanical Loading But Are Depleted from the Bone Surface During Rapid Bone Formation. JBMR Plus 6, e10593 (2022).

29. Maurel, D.B. et al. Characterization of a novel murine Sost ER(T2) Cre model targeting osteocytes. Bone Res 7, 6 (2019).

30. Wang, J.S. & Wein, M.N. Pathways Controlling Formation and Maintenance of the Osteocyte Dendrite Network. Curr Osteoporos Rep 20, 493–504 (2022).

31. Youlten, S.E. et al. Osteocyte transcriptome mapping identifies a molecular landscape controlling skeletal homeostasis and susceptibility to skeletal disease. Nat Commun 12, 2444 (2021).

32. Schurman, C.A. et al. Molecular and Cellular Crosstalk between Bone and Brain: Accessing Bidirectional Neural and Musculoskeletal Signaling during Aging and Disease. J Bone Metab 30, 1–29 (2023).

33. Raouf, A. et al. Lumican is a major proteoglycan component of the bone matrix. Matrix Biol 21, 361–367 (2002).

34. Wendel, M., Sommarin, Y. & Heinegard, D. Bone matrix proteins: isolation and characterization of a novel cell-binding keratan sulfate proteoglycan (osteoadherin) from bovine bone. J Cell Biol 141, 839–847 (1998).

35. Tashima, T., Nagatoishi, S., Sagara, H., Ohnuma, S. & Tsumoto, K. Osteomodulin regulates diameter and alters shape of collagen fibrils. Biochem Biophys Res Commun 463, 292–296 (2015).

36. Creecy, A., Damrath, J.G. & Wallace, J.M. Control of Bone Matrix Properties by Osteocytes. Front Endocrinol (Lausanne) 11, 578477 (2020).

37. Staines, K.A. et al. Hypomorphic conditional deletion of E11/Podoplanin reveals a role in osteocyte dendrite elongation. J Cell Physiol 232, 3006–3019 (2017).

38. Atkins, G.J. et al. Sclerostin is a locally acting regulator of late-osteoblast/preosteocyte differentiation and regulates mineralization through a MEPE-ASARM-dependent mechanism. J Bone Miner Res 26, 1425–1436 (2011).

39. Zhao, W. et al. Osteomodulin deficiency in mice causes a specific reduction of transversal cortical bone size. J Bone Miner Res 39, 1025–1041 (2024).

40. Zhao, W., Byrne, M.H., Wang, Y. & Krane, S.M. Osteocyte and osteoblast apoptosis and excessive bone deposition accompany failure of collagenase cleavage of collagen. J Clin Invest 106, 941–949 (2000).

41. Bivi, N. et al. Cell autonomous requirement of connexin 43 for osteocyte survival: consequences for endocortical resorption and periosteal bone formation. J Bone Miner Res 27, 374–389 (2012).

42. Mullen, C.A., Haugh, M.G., Schaffler, M.B., Majeska, R.J. & McNamara, L.M. Osteocyte differentiation is regulated by extracellular matrix stiffness and intercellular separation. J Mech Behav Biomed Mater 28, 183–194 (2013).

43. Wysolmerski, J.J. Osteocytes remove and replace perilacunar mineral during reproductive cycles. Bone 54, 230–236 (2013).

44. Powell, W.F., Jr., et al. Targeted ablation of the PTH/PTHrP receptor in osteocytes impairs bone structure and homeostatic calcemic responses. J Endocrinol 209, 21–32 (2011).

45. Yu, K. et al. Conditional inactivation of FGF receptor 2 reveals an essential role for FGF signaling in the regulation of osteoblast function and bone growth. Development 130, 3063–3074 (2003).

46. Madisen, L. et al. A robust and high-throughput Cre reporting and characterization system for the whole mouse brain. Nat Neurosci 13, 133–140 (2010).

47. Kusuzaki, K. et al. A staining method for bone canaliculi. Acta Orthop Scand 66, 166–168 (1995).

48. Dole, N.S., Yee, C.S., Schurman, C.A., Dallas, S.L. & Alliston, T. Assessment of Osteocytes: Techniques for Studying Morphological and Molecular Changes Associated with Perilacunar/Canalicular Remodeling of the Bone Matrix. Methods Mol Biol 2230, 303–323 (2021).

49. Reynolds, E.S. The use of lead citrate at high pH as an electron-opaque stain in electron microscopy. J Cell Biol 17, 208–212 (1963).

50. Yang, Y. et al. A modified tape transfer approach for rapidly preparing high-quality cryosections of undecalcified adult rodent bones. J Orthop Translat 26, 92–100 (2021).

51. Mazur, C.M. et al. Genome-wide CRISPR interference screen identifies Clip2 as a novel regulator of osteocyte maturation and morphology. bioRxiv (2025).

52. Carpenter, A.E. et al. CellProfiler: image analysis software for identifying and quantifying cell phenotypes. Genome Biol 7, R100 (2006).

53. Patro, R., Duggal, G., Love, M.I., Irizarry, R.A. & Kingsford, C. Salmon provides fast and bias-aware quantification of transcript expression. Nat Methods 14, 417–419 (2017).

54. Love, M.I., Huber, W. & Anders, S. Moderated estimation of fold change and dispersion for RNA-seq data with DESeq2. Genome Biol 15, 550 (2014).

55. Hulsen, T., de Vlieg, J. & Alkema, W. BioVenn - a web application for the comparison and visualization of biological lists using area-proportional Venn diagrams. BMC Genomics 9, 488 (2008).

