## Supplemental Legends and Methods for "Fibroblast Growth Factor Receptor Signaling in Maturing Osteoblasts Controls Cell-Matrix Interactions Critical for Osteocyte Survival"

### Supplementary Figure Legends

**Supplementary Figure 1. The zone of apoptosis comprises TUNEL+ cells closer to the endosteum rather than the periosteum.**

(**a-c**) FGFR1-CKO mice were exposed to tamoxifen from 12-16 weeks of age and tibiae were harvested for TUNEL assay (green) (**a**), immunofluorescence (IF) for OCN (**b**) and PDPN (**c**). Arrows indicate the almost identical location of TUNEL+ (**a**), OCN (**b**) and PDPN (**c**) expressing osteocytes in FGFR1-CKO tibia in the proximal (just distal to the trabecular bone), midshaft and distal tibial cortical bone. Blue, DAPI-stained nuclei; tb, trabecular bone; ct, cortical bone; bm, bone marrow. Scale bars (all panels), 100 µm.

**Supplementary Figure 2. *Fgfr2* inactivation in the DMP1-CreER lineage does not result in osteocyte apoptosis.**

Twelve-week-old Control *Fgfr2*$^{f/f}$ and FGFR2-CKO (Dmp1-CreER; *Fgfr2*$^{f/f}$) mice were exposed to tamoxifen for 4 weeks to induce DMP1-CreER and analyzed at 16 weeks of age (also see Fig. 1b). (**a, b**) H&E staining of the tibial cortical bone from Control *Fgfr2*$^{f/f}$ (**a**) and FGFR2-CKO (**b**) mice showing the presence of viable nuclei (hematoxylin stained, blue) throughout the cortical bone. (**c-e**) TUNEL assay on the tibiae from Control *Fgfr2*$^{f/f}$ (**c**) and FGFR2-CKO (**d, e**) mice showing the absence of TUNEL+ osteocytes in the tibial diaphyseal bone (**d**), but normal developmental apoptosis in the hypertrophic zone of the growth plate (**e**). Blue, DAPI-stained nuclei; tb, trabecular bone; ct, cortical bone; bm, bone marrow; mu, muscle; gp, growth plate. Scale bars (all panels), 100 µm.

**Supplementary Figure 3. Immunofluorescence (IF) analysis of osteoblast markers.**

(**a-c**) Mice were exposed to tamoxifen from 12-16 weeks of age and tibiae were harvested for IF analysis for periostin (POSTN) (**a**) in *Fgfr1*$^{f/f}$ and FGFR1-CKO tibiae, PDPN (**b**) and Alkaline Phosphatase (ALPL) (**c**) in DFF and FGFR1,2-DCKO tibiae. Arrows in (**a**) indicate expression of POSTN in the periosteum of both *Fgfr1*$^{f/f}$ and FGFR1-CKO tibiae. Arrows in (**b**, **c**)indicate the expression of PDPN (**b**) and ALPL (**c**) in FGFR1,2-DCKO tibia. Blue; DAPI-stained nuclei; bm, bone marrow; ct, cortical bone. Scale bars (all panels) 100 µm.

**Supplementary Figure 4. Colocalization of TUNEL+ osteocytes with OCN and ALPL.**

FGFR1,2-DCKO mice were exposed to tamoxifen from 12-16 weeks of age and tibiae were harvested at 16 weeks of age. TUNEL assay (green) and IF were done on the same histological sections.
(**a**) TUNEL assay (green) and IF for OCN (red). The dashed square identifies the region enlarged in (**a**) with signals from TUNEL staining (**b**, green) and OCN staining (**c**, red) shown individually. Arrows indicate cells that are both TUNEL+ and express OCN.
(**d**) TUNEL assay (green) and IF for ALPL (red). The dashed square identifies the region enlarged in (**d**) with signals from TUNEL staining (**e**, green) and ALPL staining (**f**, red) shown individually. Arrows indicate cells that are both TUNEL+ and express ALPL.
Scale bars: (**a**) 100 µm; (**b, c, d**) 50 µm; (**e, f**) 10 µm.

**Supplementary Figure 5. Loss of the LCN in FGFR1-CKO tibia.**

FGFR1-CKO mice were exposed to tamoxifen from 12-16 weeks of age and tibiae were harvested at 16 weeks of age. Silver staining (brown) of Control *Fgfr1*$^{f/f}$ (**a, b**) and FGFR1-CKO (**d, e**) midshaft and distal tibia. TUNEL staining (green) (**c, f**) is shown for positional correlation showing the lack of LCN in regions of TUNEL+ osteocytes in FGFR1-CKO tibia (**f**). Arrows indicate the edge of the intact LCN within the cortical bone and show a severe reduction in LCN staining between the arrow to the endosteum in FGFR1-CKO tibiae. Blue; DAPI-stained nuclei; bm, bone marrow; ct, cortical bone. Scale bars (all panels): 100 µm.

**Supplementary Figure 6. Measurement of dendrite length and proliferation assays with Ocy454 cells.**

(**a**) Quantification of dendrite length (edge of nucleus to end of projection) in Ocy454 osteocyte-like cells treated for 48 hr with FGF2 (5ng/mL) with or without BGJ398 (200 nM) showing decreased dendrite length, comparing treatment with FGF2 and inhibition of FGFR signaling by BGJ398.
(**b**) Quantification of proliferation in Ocy454 osteocyte-like cells treated for 48 hr with FGF2 (5ng/mL) with or without BGJ398 (200 nM).
ANOVA with Dunn's multiple comparisons test, * $P$<0.05, ** $P$<0.01, *** $P$<0.001, **** $P$<0.0001.

**Supplementary Figure 7. Dynamic histomorphometry of Control and FGFR deficient cortical bone.**

(**a**) Experimental plan for tamoxifen-induced *Fgfr1* inactivation and alizarin red and calcein green administration.
(**b**) Representative images of alizarin red and calcein green incorporation in the tibia of *Fgfr1*$^{f/f}$ and FGFR1-CKO mice after four weeks of tamoxifen administration showing appositional bone growth only on the endosteal side of the tibial cortical bone. The arrows indicate negligible incorporation of alizarin and calcein on the periosteal side. The dashed square represents regions magnified below. Besides alizarin red, the FGFR1-CKO mice also exhibit red signals from the *ROSA*$^{TDT}$ reporter following tamoxifen induction.
(**c**) Analysis of bone formation rate/bone surface (BFR/BS) and mineralizing surface/bone surface (MS/BS) in the tibia from FGFR1-CKO (CKO) and FGFR1,2-DCKO (DCKO) mice compared to the tibia from Control (Ctrl, *Fgfr1*$^{f/f}$ or DFF) mice. Data shown as mean ± SD. One way ANOVA with multiple comparisons (Tukey), n=4-5, *$P$<0.05, **$P$<0.01; ns, not significant.

**Supplementary Figure 8. Osteocyte death is observed only in the ECM of cortical bone formed after *Fgfr1* inactivation.**

(**a**) *Fgfr1,2*$^{f/f}$ (DFF) and FGFR1,2-DCKO mice, which do not harbor the *ROSA*$^{TDT}$ reporter allele, were exposed to tamoxifen from 12-16 weeks of age to induce DMP1-CreER. Mice were injected with alizarin red at 12 weeks of age, to mark the boundary of the newly formed bone (arrow). Tibiae were harvested and assessed for apoptosis by TUNEL staining (green) (also see Fig. 5**a**, **b**).
(**b**) Fourteen-week-old *Fgfr1,2*$^{f/f}$ (DFF) and FGFR1,2-DCKO mice, which do not harbor the *ROSA*$^{TDT}$ reporter were subjected to mechanical loading to induce bone on the periosteal side of the posterior tibia (also see Fig. 5**c**, **d**). Mice were injected with alizarin red on days 4 and 5 of loading to mark newly formed bone. TUNEL staining of the loaded tibia showed apoptosis (green) only within the region of newly formed bone between the periosteum (arrowhead) and the alizarin red label. Arrow indicates the boundary of the newly formed bone within which apoptosis is observed.
Blue, DAPI-stained nuclei; ct, cortical bone; bm, bone marrow; mu, muscle.

**Supplementary Figure 9. Inactivation of *Fgfr1* with Sost-CreER does not affect osteocyte viability.**

Sost-CreER; *ROSA*$^{TDT}$ and SOST-FGFR1-CKO mice were exposed to tamoxifen from 12 to 16 weeks of age and analyzed at 16 weeks of age.
(**a**) IF staining of femoral cortical bone for TDT showing expression of the *ROSA*$^{TDT}$ reporter in osteocytes, some periosteal fibroblasts and muscle cells from Sost-CreER; *ROSA*$^{TDT}$ mice.
(**b**) qRT-PCR analysis showing reduction of *Fgfr1* expression in mRNA harvested from flushed cortical hindlimbs of SOST-FGFR1-CKO mice. Data shown as mean ± SD. Unpaired *t*-test, n=5, ***$P$<0.001.
(**c**) Representative images showing the absence of TUNEL+ osteocytes in the tibial cortical bone of SOST-FGFR1-CKO mice.
Blue; DAPI-stained nuclei; bm, bone marrow; ct, cortical bone. Scale bar (all panels): 100 μm.

**Supplementary Figure 10. Analysis of cortical bone gene expression and RNA-seq data.**

(**a-b**) Analysis of the read count from exons 2-17 of the *Fgfr1* coding region from the RNA harvested from flushed cortical bones in 13 mRNA samples (from 6 *Fgfr1*$^{f/f}$ and 7 FGFR1-CKO mice) submitted for bulk RNA sequencing. Four Control samples expressing the highest levels of *Fgfr1* and four FGFR1-CKO samples expressing the lowest levels of *Fgfr1* were selected for further analysis. (**a**) Each dot represents the read count of an exon from 2-17 of *Fgfr1* for each sample. Samples are rank ordered by average read count. (**b**) Relative expression based on multiple comparisons from ANOVA analysis of the data in panel (**a**). See Methods for details.
(**c**) qRT-PCR analysis of DFF Control and FGFR1,2-DCKO diaphyseal RNA for *Col1a1*, *Sp7*, *Gja1* and *Pdpn* validating the differential down-regulation of these genes identified by RNA-seq DEG analysis. Data shown as the mean ± SD. Unpaired *t*-test, n=5, *$P$<0.0, **$P$<0.01.
(**d, e**) Heat map showing relative expression from mouse mRNA seq data of genes identified as associated with ECM regulation (**d**) and Neuronal regulation (**e**) from pathway enrichment analysis of Control (*Fgfr1*$^{f/f}$) and FGFR1-CKO samples. Changes in orange indicate increased relative expression, and blue indicates decreased relative expression.

**Supplementary Figure 11. Heatmaps comparing cell stress and ECM production in mouse and Ocy454 RNA-seq data.**

(**a**) Heat map from mouse mRNA seq data showing increased relative expression of genes identified as associated with cellular stress from pathway enrichment analysis of Control (*Fgfr1*$^{f/f}$) and FGFR1-CKO samples.
(**b**) Heat map from Ocy454 mRNA seq data showing lack of regulation of cell stress genes identified in the mouse data.
(**c-d**) Heat map showing relative expression from mouse mRNA seq data of 24 common ECM associated genes (**c**) and 15 common neuronal regulatory genes (**d**) shared between the *in vivo* mouse model of *Fgfr1* inactivation and *in vitro* Ocy454 manipulation by FGF2 and FGFR inhibitor, BGJ398. Changes in orange indicate increased relative expression, and blue indicates decreased relative expression.

**Supplementary Figure 12. Regulation of apoptosis gene expression in mouse and Ocy454 RNA-seq data.**

(**a**) Heat map from mouse mRNA seq data showing differential expression of 26 genes identified as associated with apoptosis.
(**b**) Heat map from Ocy454 mRNA seq data comparing differential gene expression with the apoptosis genes identified in mouse. Asterisks indicate genes differentially expressed in both mouse and Ocy454 data.
(**c, d**) Comparison of changes in gene expression of the 26 apoptosis-related mouse DEGs in mouse and Ocy454 RNA seq data. (**c**) genes upregulated in CKO mouse tend to be downregulated in BGJ398-treated Ocy454 cells (black dots). (**d**) genes downregulated in CKO mouse tend to be upregulated in BGJ398-treated Ocy454 cells (black dots). Grey dots, not significantly changed. Wilcoxon matched-pairs signed rank test, upregulated genes (*$P$<0.02), downregulated genes (***$P$<0.001).

**Supplementary Figure 13. *Fgfr1* inactivation in the DMP1-CreER lineage results in downregulation of bone mineralization genes.**
(**a**) Heat map from mouse mRNA seq data showing differential expression of 47 genes identified as associated with bone mineralization from pathway enrichment analysis.
(**b**) RPM (reads per million) for *Fgfr1*, *DMP1*, *Phex*, *Enpp1* and *Fgf23* from Control (*Fgfr1*$^{f/f}$) and FGFR1-CKO mice showing their down-regulation in mRNA seq data.
(**c**) qRT-PCR analysis of Control (*Fgfr1*$^{f/f}$) and FGFR1-CKO diaphyseal RNA for *Dmp1*, *Phex*, *Enpp1* and *Fgf23* validating the differential down-regulation of these genes identified by RNA-seq DEG analysis in (**b**).

**Supplementary Figure 14. Abnormal ECM in regions of new bone formation following FGFR ablation after four weeks of tamoxifen induction.**

Mice were exposed to tamoxifen from 12-16 weeks of age and tibiae were harvested for ultrastructural analysis.
(**a, b**) Scanning electron microscopy (SEM) images of *Fgfr1*$^{f/f}$ and FGFR1-CKO tibiae in regions of new endosteal bone formation under different magnifications. Arrows point to the organized lamellar banding pattern in *Fgfr1*$^{f/f}$ bone, and to the irregular banding pattern in FGFR1-CKO ECM of the collagen fibrils surrounding dead osteocytes (*). Dashed squares in (**b)** show the area magnified in the inset.
(**c-d**) Transmission electron microscopy (TEM) images of *Fgfr1*$^{f/f}$ and FGFR1-CKO tibiae in

regions of new endosteal bone formation under different magnifications. The ECM of FGFR1-CKO bone shows irregular ridges surrounding dead osteocytes (*). Dashed squares in (**c**) are shown magnified in (**d**).
blc, bone lining cell; bm, bone marrow; ct, cortical bone. Scale bars: (**a**) 5 µm; (**b**) 500 nm; insets in (**b**), 250 nm; (**c**) 10 µm; (**d**), 2 µm.

**Supplementary Methods:**

**Ocy454 viability and proliferation assays**

FGFR1/2/3 inhibitor BGJ398 (Selleck Chemical) was reconstituted in DMSO at 1 mM and diluted 1:100 in media to create a working solution of 10 µM. DMSO was diluted 1:100 in media to create a working solution of vehicle. Working solutions were diluted 1:50 into media for a final treatment dose of 200 nM BGJ398. FGF2 (Proteintech Group # 50-197-3960) was reconstituted in PBS + 0.1% BSA at 100 µg/mL and then diluted into media for a final treatment dose of 5 ng/mL. All experiments were performed in complete media (alpha-MEM supplemented with 10% FBS and 1% antibiotic/antimycotic).

Ocy454 cells were plated onto plastic tissue culture-treated plates in complete media and cultured at 33°C. Starting the next day, cells were pretreated for one hour with FGFR inhibitor BGJ398 or DMSO vehicle. Treatment media was then applied in each of four conditions: DMSO, DMSO + FGF2 (5 ng/mL), BGJ398, BGJ398 + FGF2 (5 ng/mL) for a total of 48 hours. After 46 hours of treatment, EdU (5-ethynyl-2´-deoxyuridine) was added to treatment media to reach a final concentration of 10 µM, and cells continued to incubate at the permissive temperature (33°C) for two hours.

At the end of treatment, cells without EdU treatment were trypsinized and collected for Annexin V/propidium iodide (PI) labeling for analysis of viability, and cells with and without EdU treatment were trypsinized and collected for EdU detection for analysis of proliferation.

For Annexin V/PI, approximately 100,000 cells per condition were immediately washed and resuspended in Annexin V Binding Buffer (BioLegend), then labeled at room temperature for 15 minutes using a FITC Annexin V Apoptosis Detection Kit with PI (BioLegend). Fluorescence was measured in an Attune NxT flow cytometer.

For EdU detection, approximately 100,000 cells per condition were washed and fixed, then permeabilized and labeled with the Click-iT Plus EdU Alexa Fluor 488 Flow Cytometry Assay Kit (Invitrogen). Fluorescence was measured in an Attune NxT flow cytometer.

Dendrite length from the edge of the nucleus to the end of the phalloidin-stained projection was measured manually for the longest dendrite of each cell using ImageJ.

**qRT-PCR assays**

For quantitative gene expression analysis, TaqMan off-the-shelf gene expression assays (Thermo Fisher Scientific) and TaqMan Fast Universal PCR Master Mix (2X) (Thermo Fisher Scientific) were used. cDNA was added to TaqMan Fast Universal Fast Master Mix with TaqMan probes and primer sets specific for the gene of interest, and assays performed as per the manufacturer's protocol. Samples were added to a 96 well plate in triplicates for both house-keeping gene (*Hprt1*) and target genes. PCR was performed using an iCycler (Life Technologies) with the manufacturer's Fast protocol and *ΔCT* values calculated using *Ct* values of Hprt1 as the house keeping control.
