## Supplemental Figures for "Fibroblast Growth Factor Receptor Signaling in Maturing Osteoblasts Controls Cell-Matrix Interactions Critical for Osteocyte Survival"

**Supplementary Figure 1**

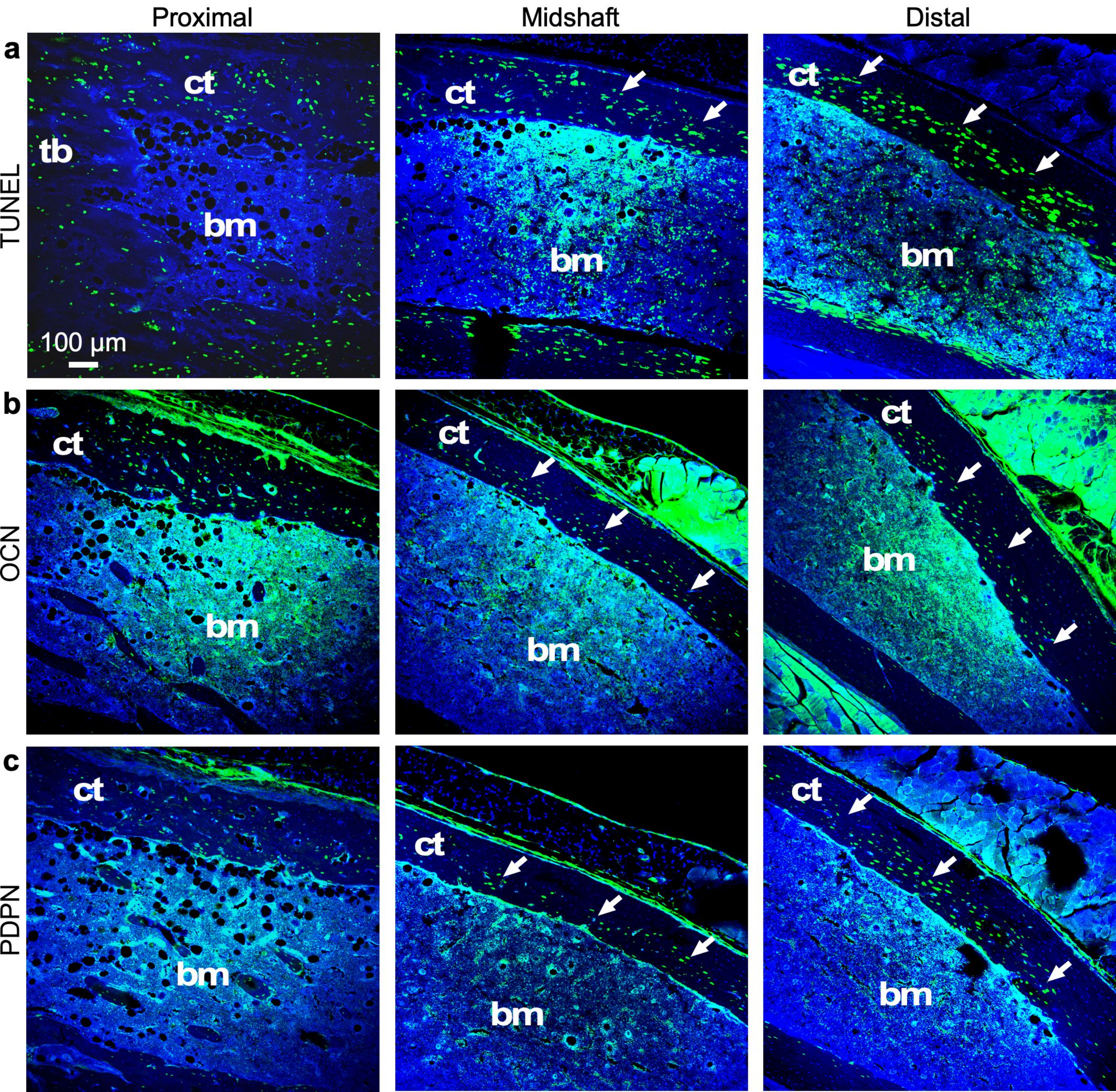

### Supplementary Figure 2

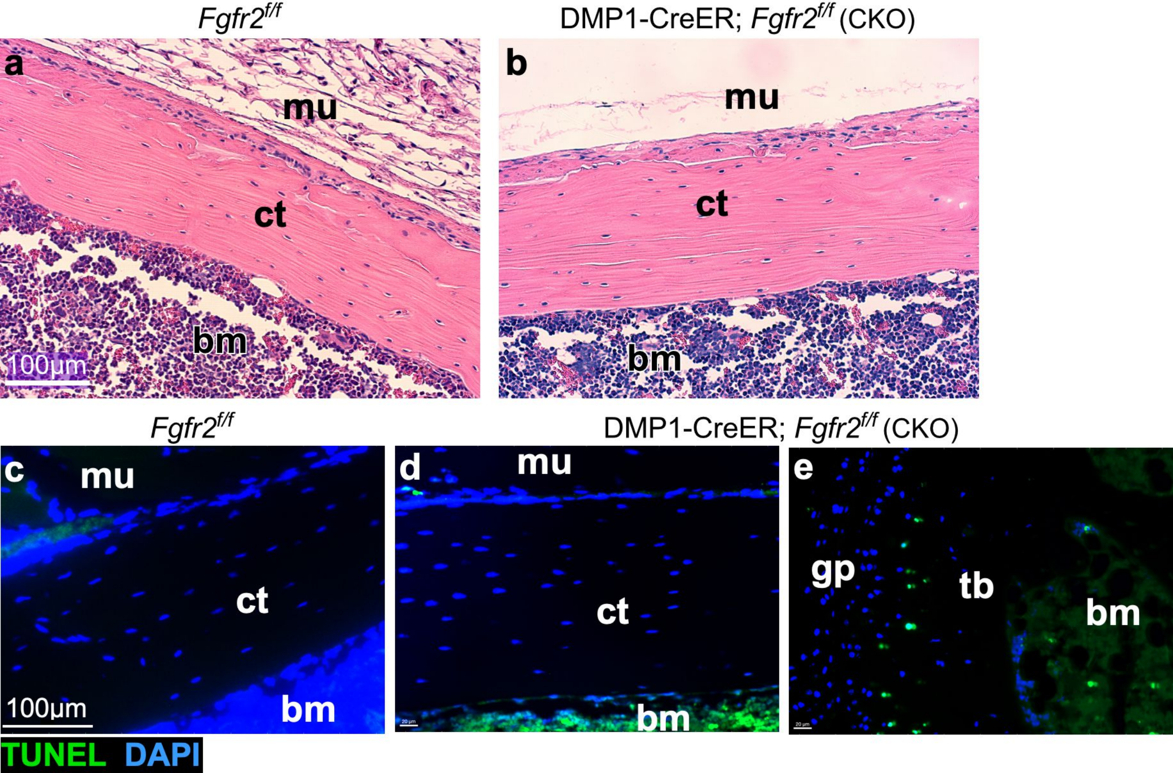

### Supplementary Figure 3

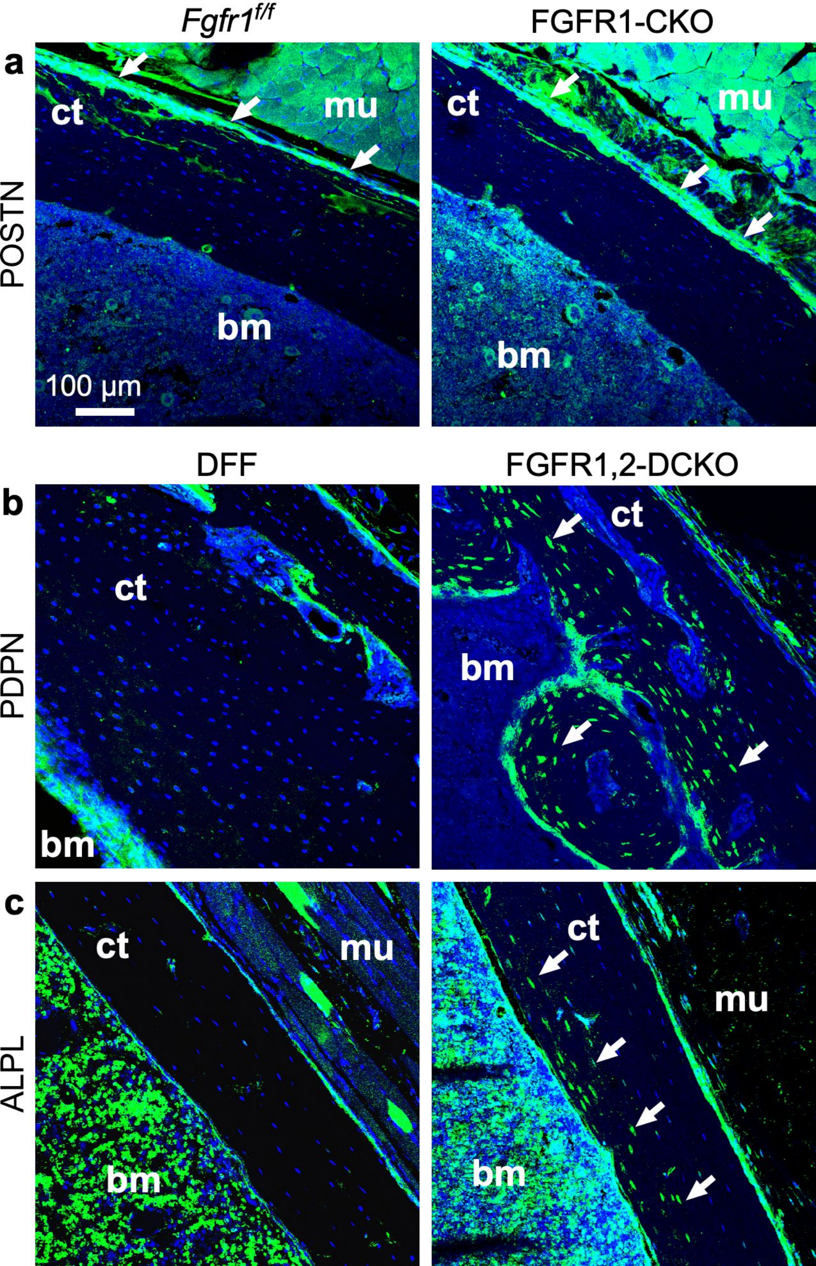

#### Supplementary Figure 4

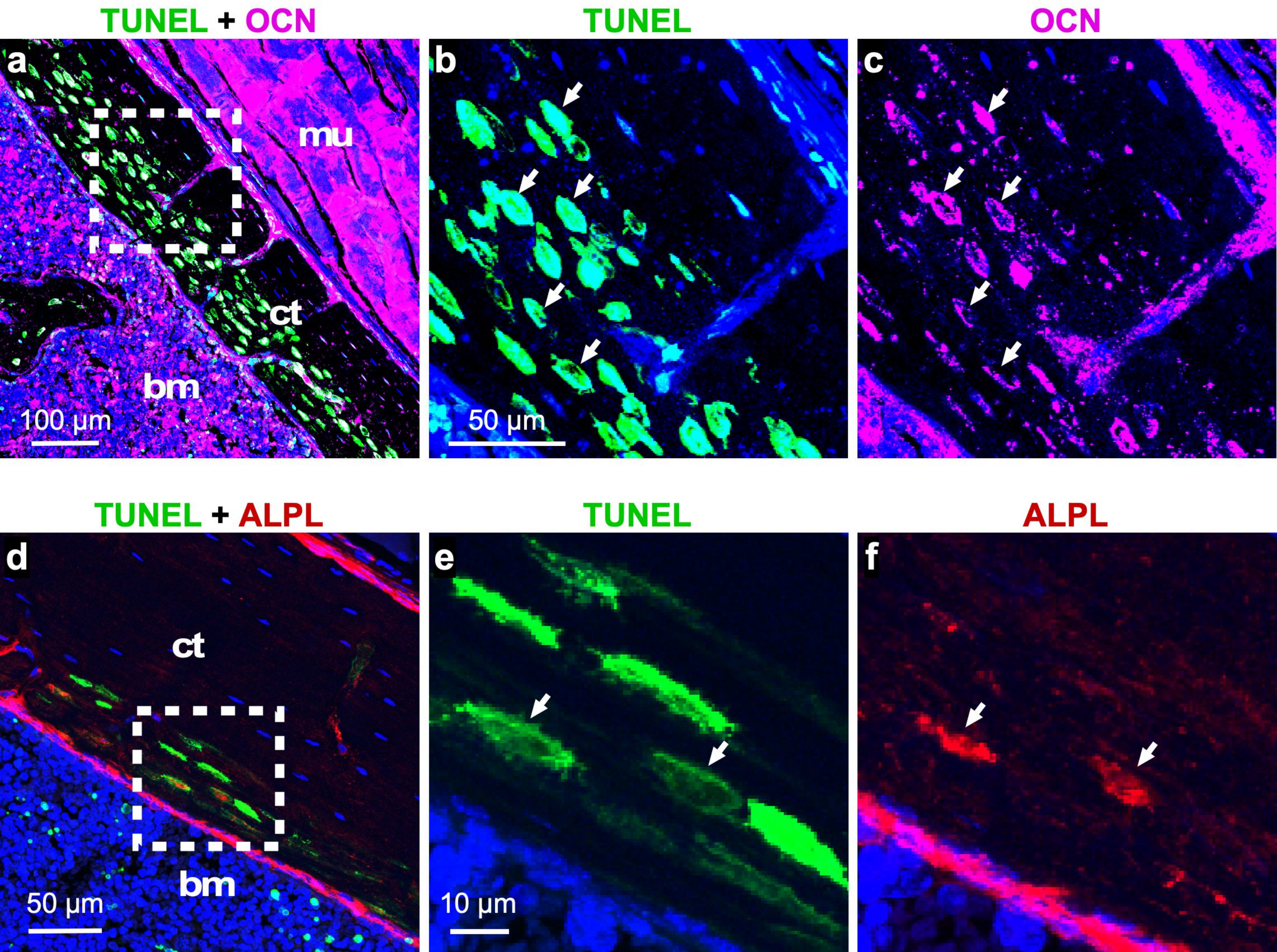

**Supplementary Figure 5**

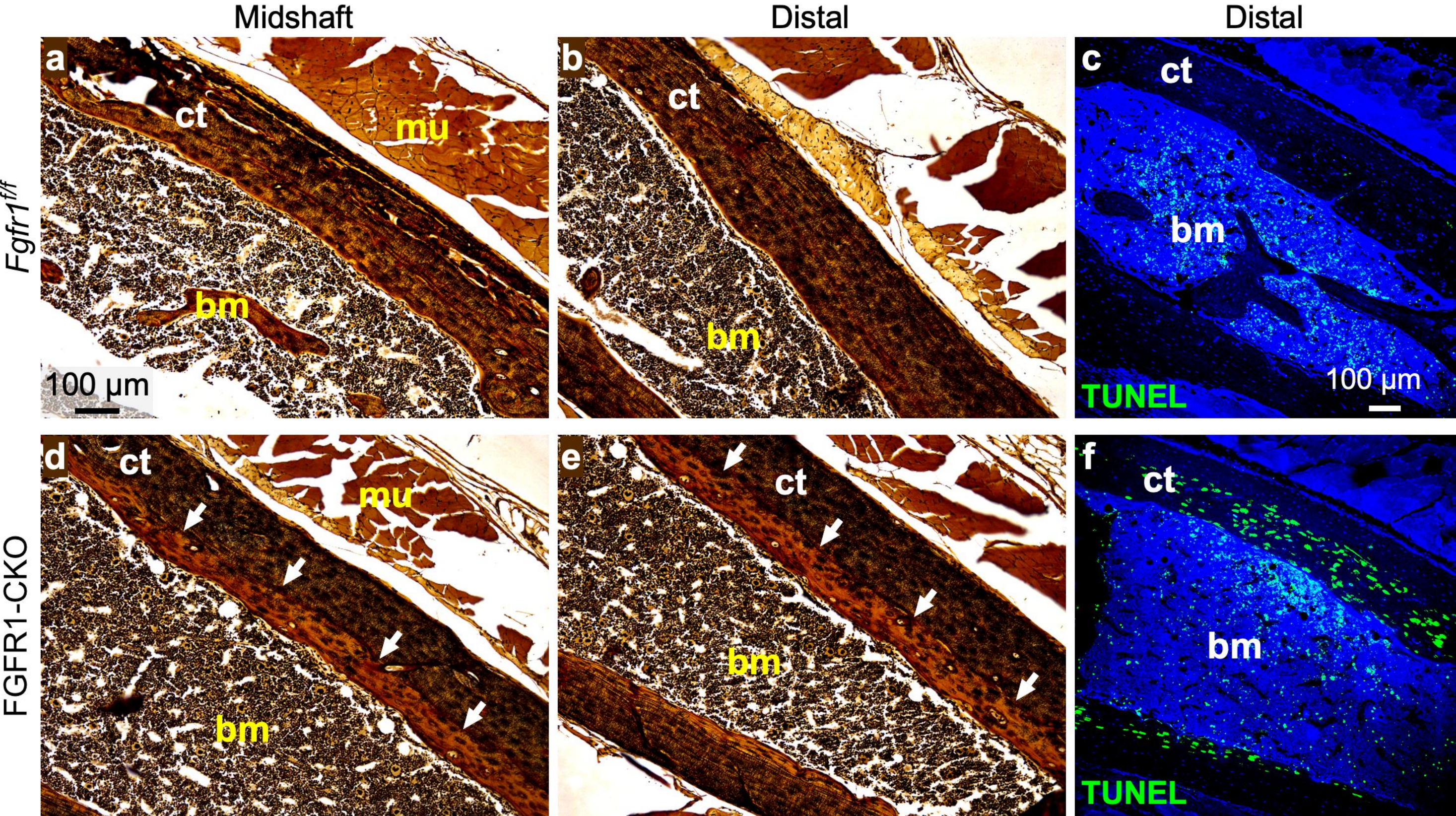

### Supplementary Figure 6

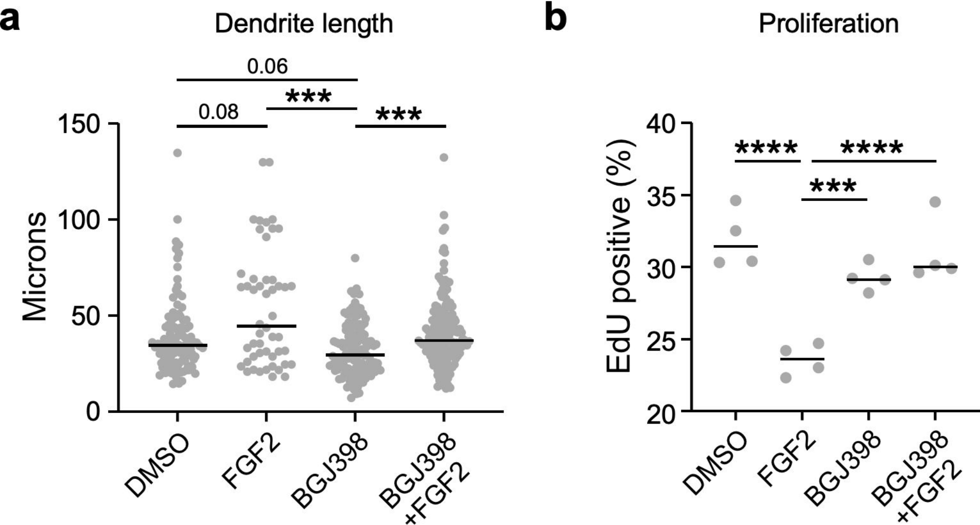

### Supplementary Figure 7

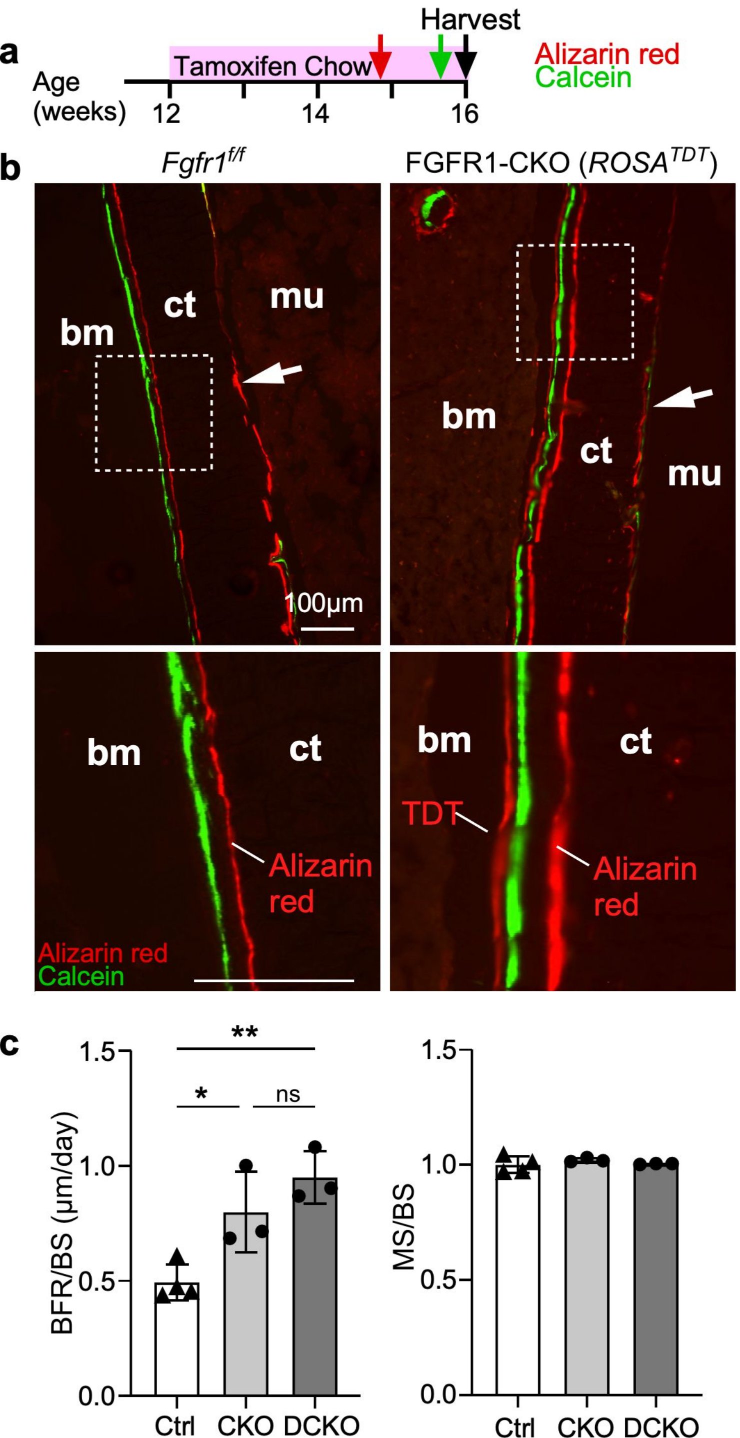

### Supplementary Figure 8

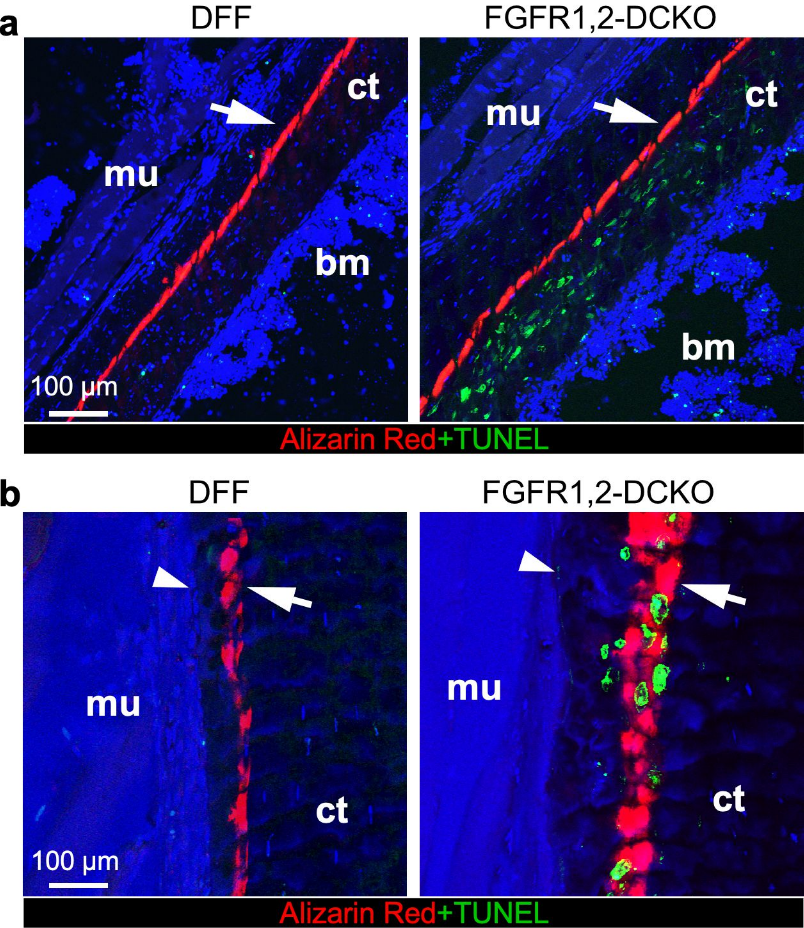

### Supplementary Figure 9

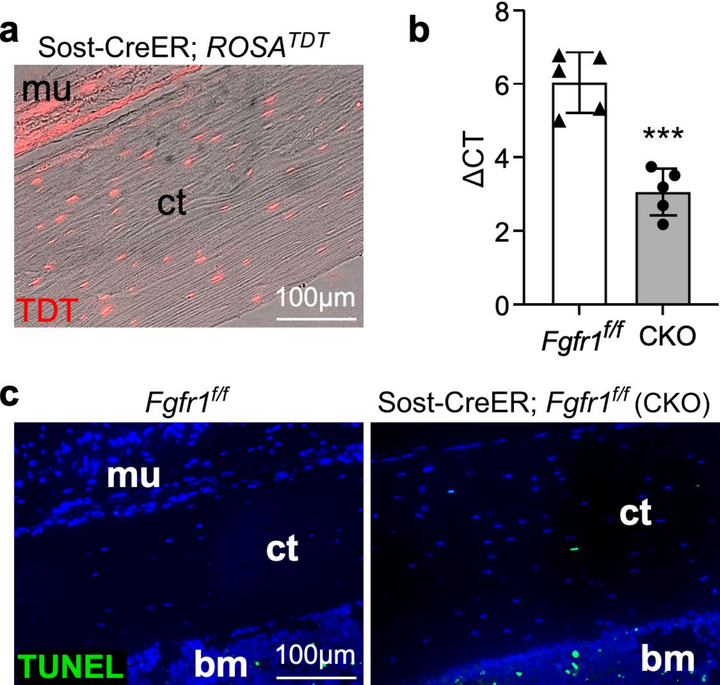

#### Supplementary Figure 10

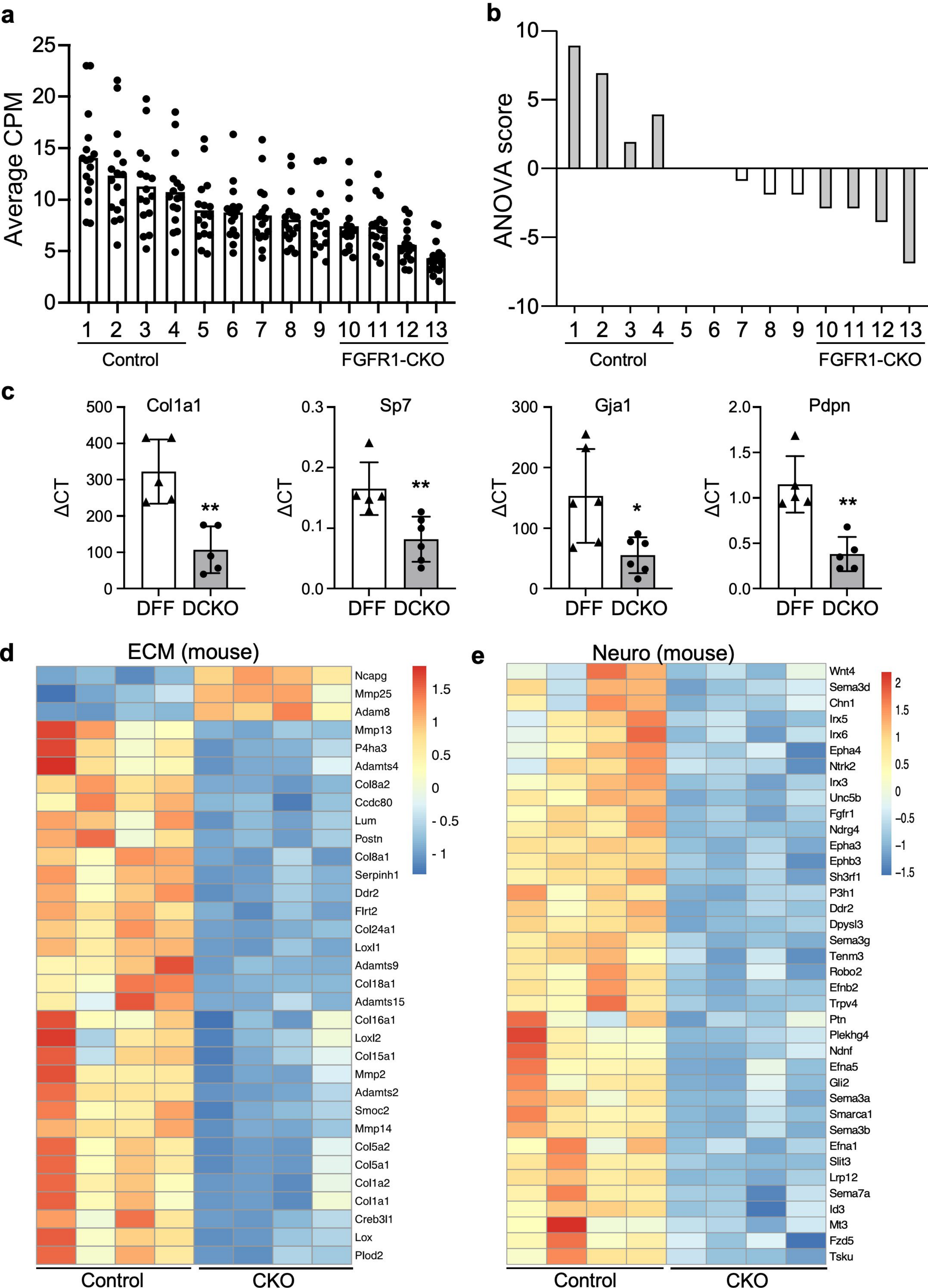

### Supplementary Figure 11

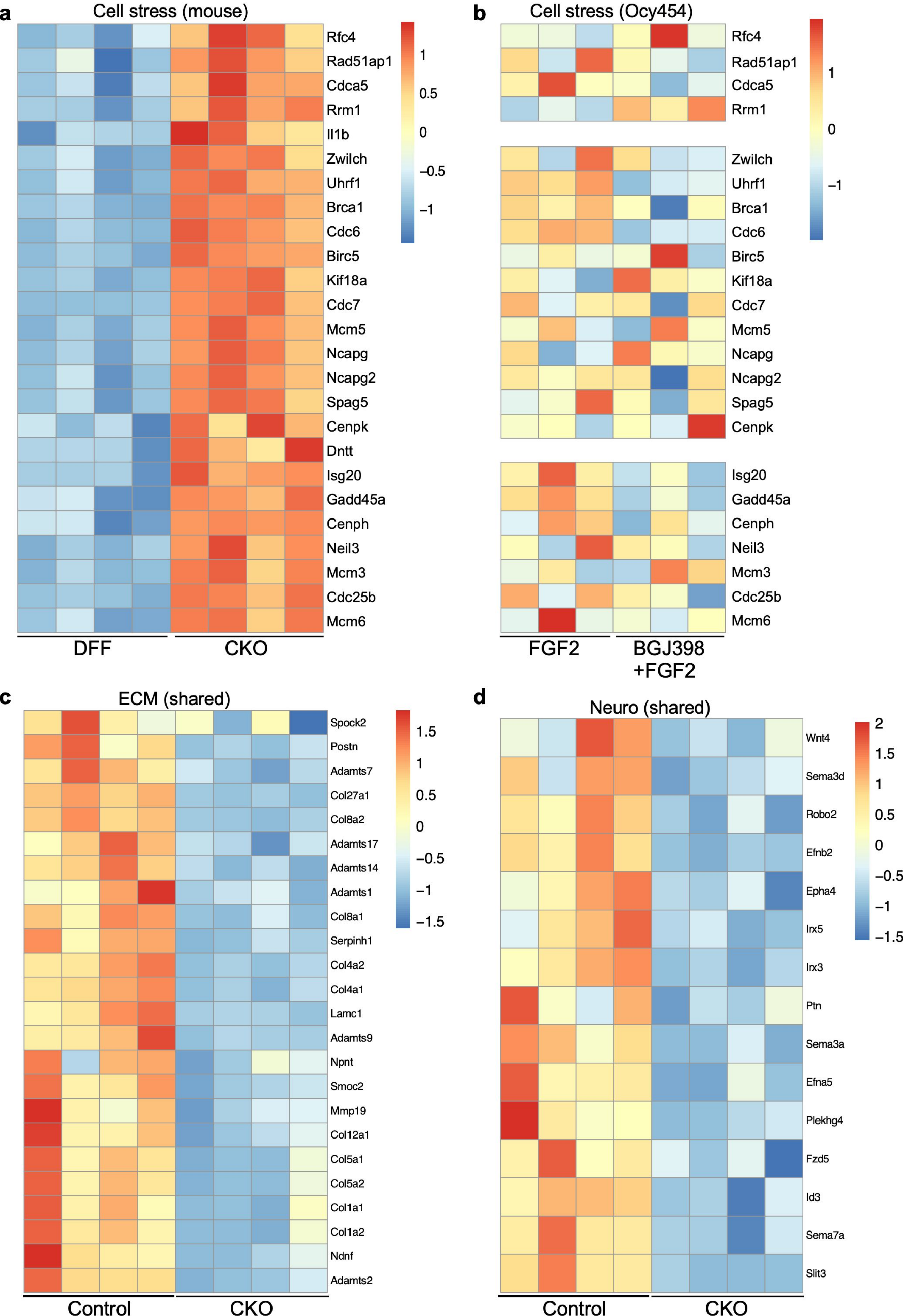

### Supplementary Figure 12

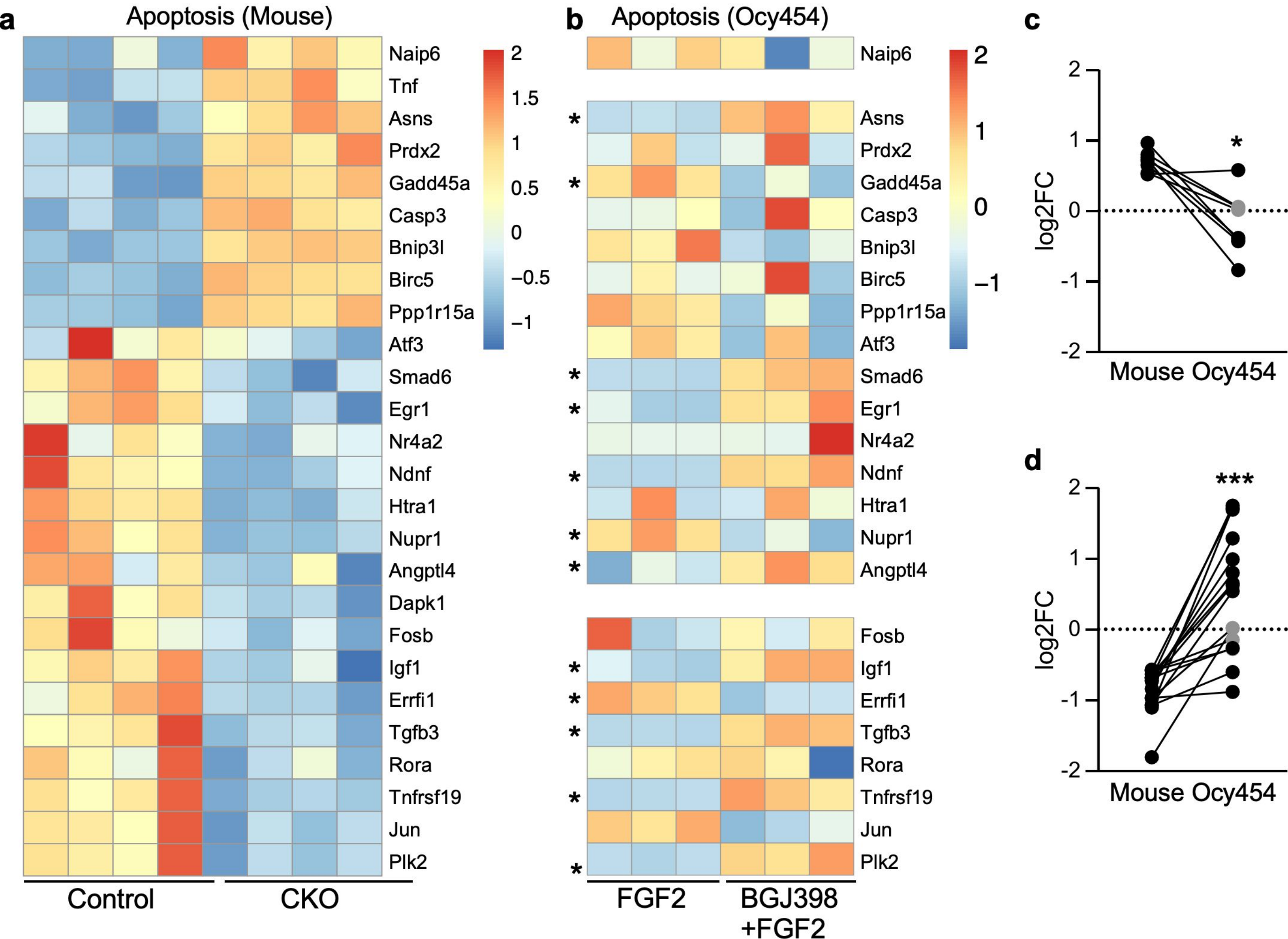

#### Supplementary Figure 13

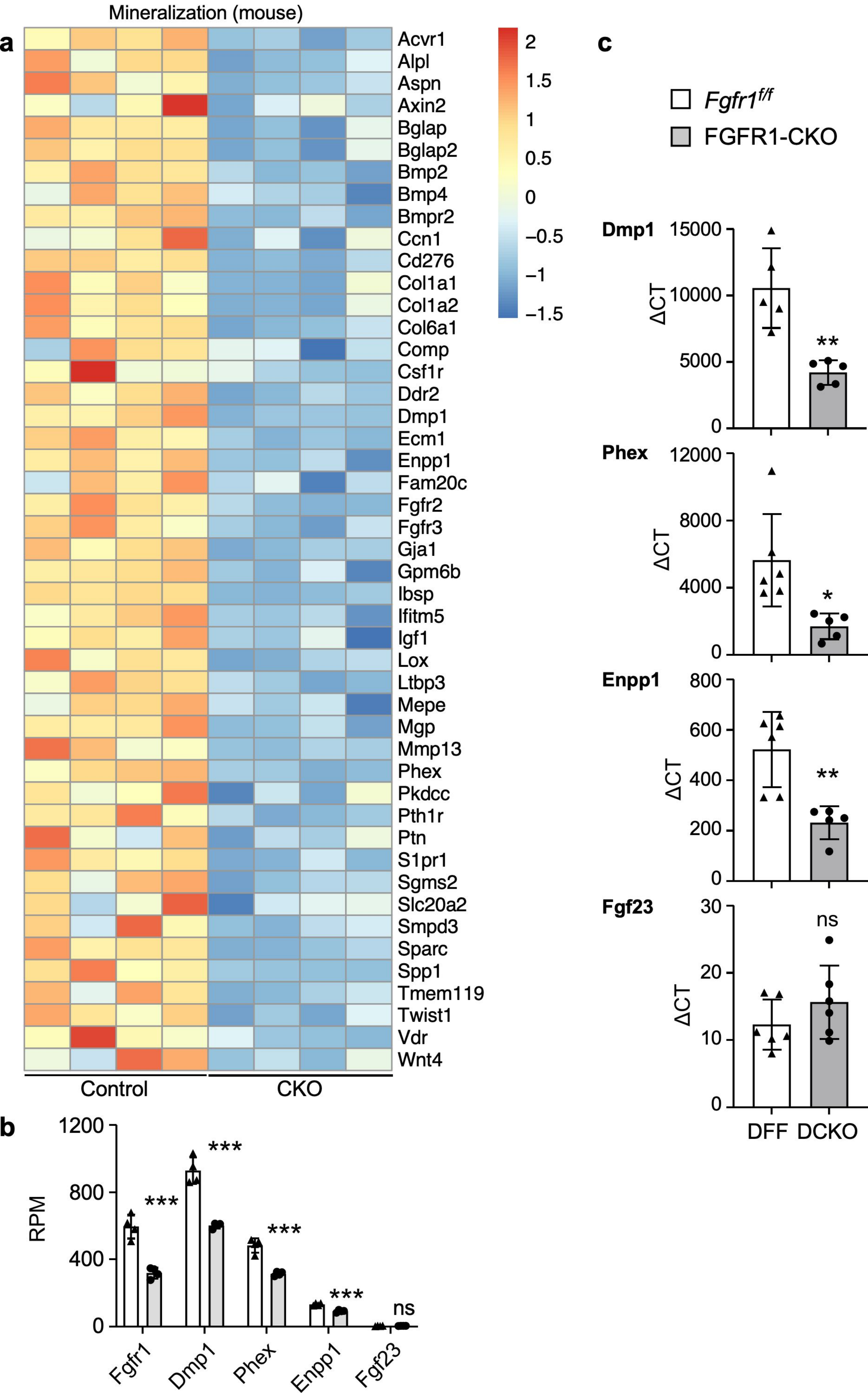

### Supplementary Figure 14

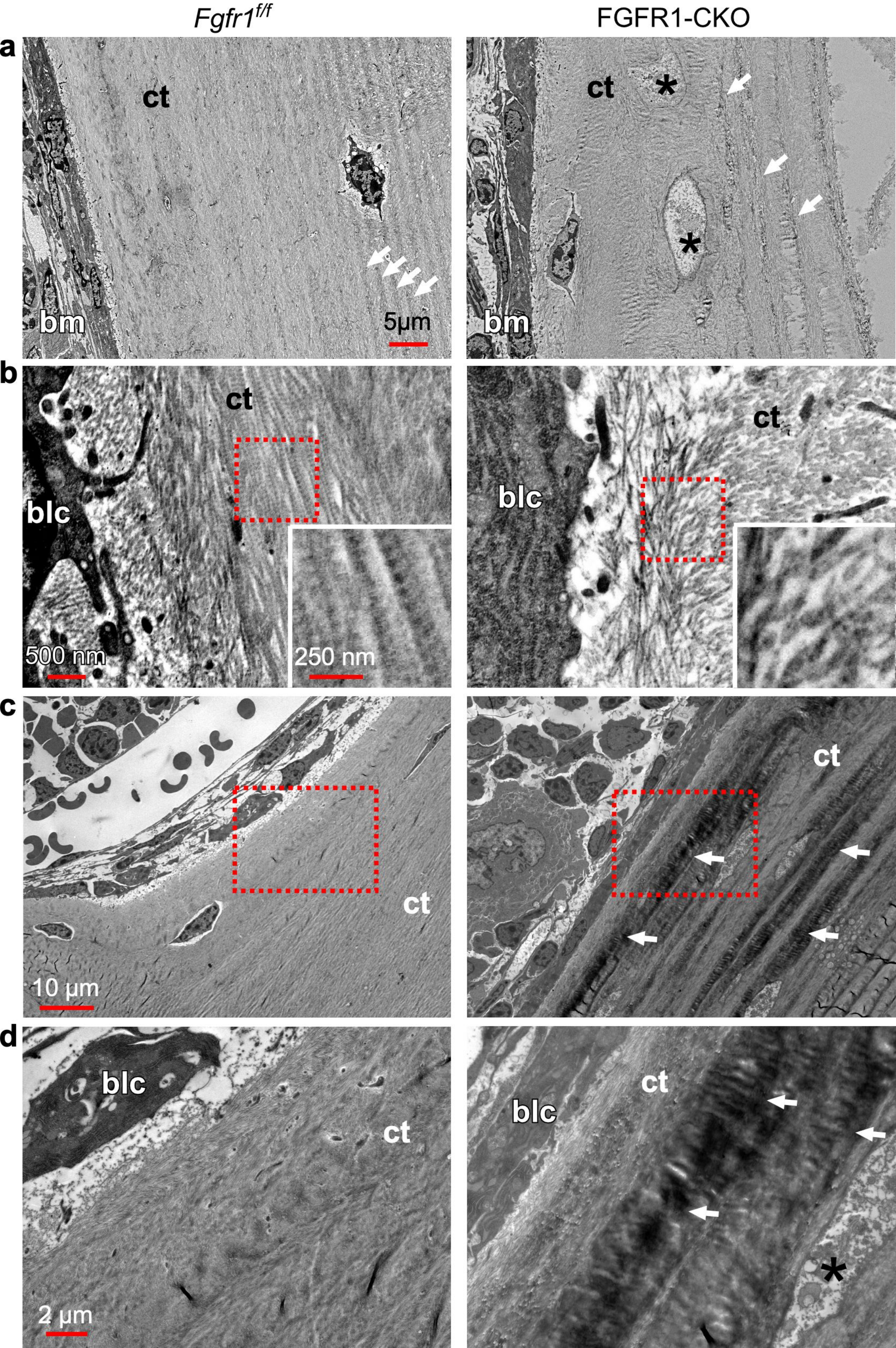
